# Molecular mechanism of dynein-dynactin activation by JIP3 and LIS1

**DOI:** 10.1101/2022.08.17.504273

**Authors:** Kashish Singh, Clinton K. Lau, Giulia Manigrasso, José B. Gama, Reto Gassmann, Andrew P. Carter

**Affiliations:** MRC Laboratory of Molecular Biology, Francis Crick Ave, Cambridge, CB2 0QH, UK; Instituto de Investigação e Inovação em Saúde – i3S / Instituto de Biologia Molecular e Celular – IBMC, Universidade do Porto, 4200-135 Porto, Portugal; Current affiliation: Department of Biochemistry, University of Oxford, Oxford, OX1 3QU, UK

## Abstract

Microtubule motors, like cytoplasmic dynein-1, are tightly regulated to prevent inappropriate activity in cells. Dynein functions as a ∼4 MDa complex containing its cofactor dynactin and a cargo-specific coiled-coil adaptor. However, how dynein and dynactin recognise diverse adaptors, how they interact with each other during complex formation, and the role of critical regulators such as LIS1 remain unclear. To address this, we determine the cryo-EM structure of dynein-dynactin on microtubules with LIS1 and the lysosomal adaptor JIP3. We show how JIP3 specifies complex formation despite a shorter coiled coil and lack of identifiable motifs compared to typical adaptors. We find LIS1 directly binds dynactin’s p150 subunit, closely tethering it to dynein. This interaction is necessary for dynein’s cellular and *in vitro* activity. We also show how dynein’s intermediate chain relieves p150’s autoinhibition to enable LIS1 binding. Together, our data suggest LIS1 and p150 constrain dynein-dynactin to ensure efficient complex formation.

## Introduction

Cytoskeletal motors play crucial roles in cellular organization, division and function^1–5^. These molecular machines are tightly regulated to ensure correct recruitment and activation^6,7^, with defects in these control mechanisms leading to developmental and neurodegenerative diseases^8–10^.

The motor cytoplasmic dynein-1 (dynein) drives the majority of movement toward the minus-ends of microtubules in our cells. It is unusual in requiring a large, 1.1 MDa, cofactor called dynactin^11^. This binds dynein in the presence of cargo-specific ‘activating adaptors’ from one of several different adaptor families^1,12,13^. The assembly of dynein-dynactin-adaptor (DDA) complexes is a key step in regulating dynein, converting it into a motor capable of long-distance movement^12–14^. In the cell, this assembly process requires the presence of additional factors^15,16^, with a crucial example being the regulator LIS1^17^. The complexity of DDA complexes raises two questions: how can their assembly be driven by such a diverse set of adaptors, and how does LIS1 stimulate this process?

Activating adaptors share a long N-terminal coiled coil and motifs for interacting with specific parts of dynein or dynactin. Dynein is a dimer of heavy chains (DHCs), intermediate chains (DICs), light intermediate chains (DLICs) and light chains^18–21^. All known activating adaptors contact the flexible C-terminal helix of DLIC (DLIC^helix^). They do this by a variety of different motifs or domains: for example, the CC1 box motif in BICD2, the Hook domain in HOOK3, and the EF-Hand domain in Rab11FIP3^1,22–24^ (Figure 1A). Some adaptors also contain a motif in their N-terminal coiled coil which binds the DHC and is referred to as the heavy chain binding site 1 (HBS1)^25–27^. Finally, many adaptors contain a Spindly motif which contacts dynactin^25,27–29^.

**Figure 1.**
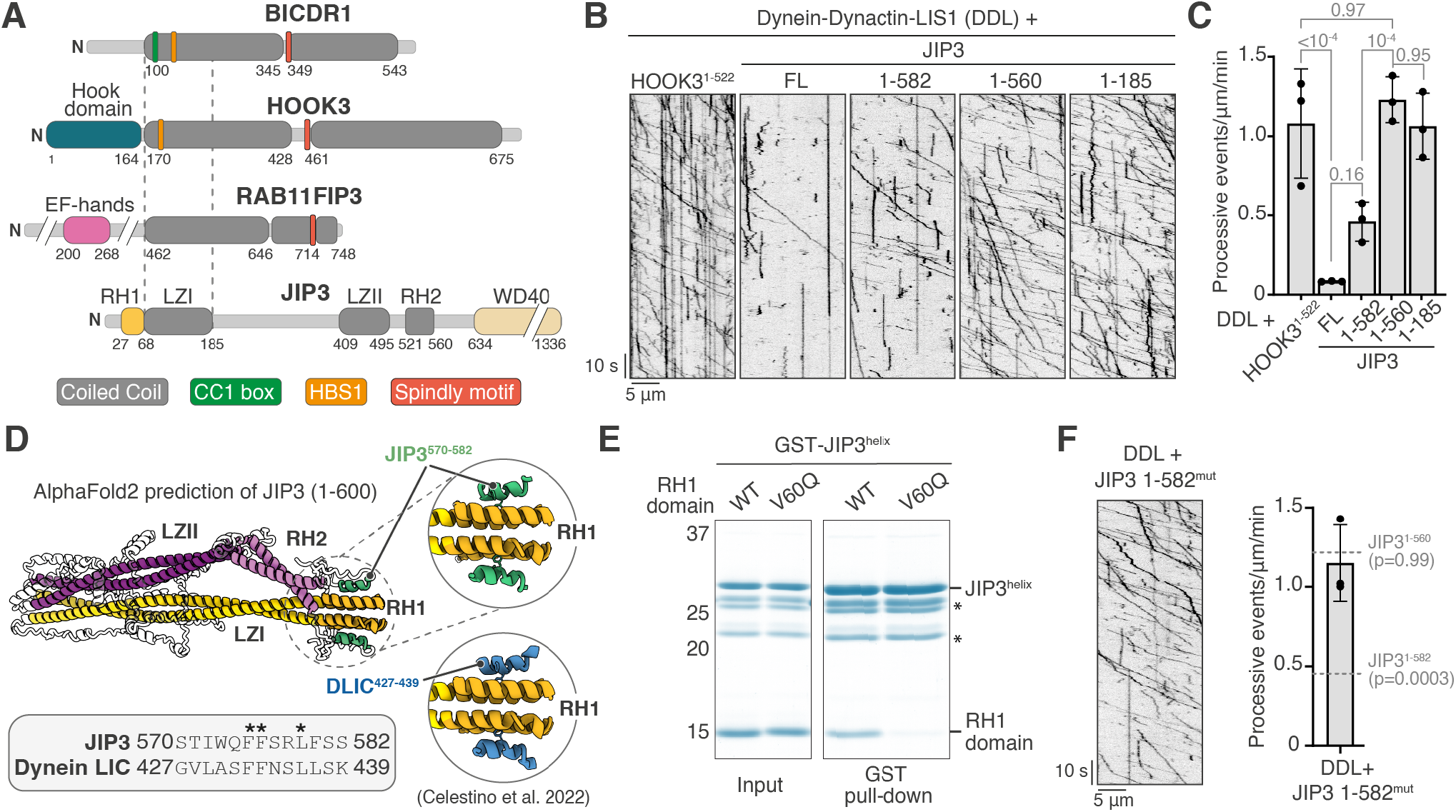
JIP3 is an autoinhibited activating adaptor for dynein motility *in vitro*. **(A)** Schematic representation of BICDR1 (also known as BICDL1), HOOK3, Rab11FIP3 and JIP3. The dotted lines illustrate the short coiled coil of JIP3 as compared to known dynein activating adaptors. **(B)** Kymographs of TMR-dynein-dynactin-LIS1 in the presence of HOOK3^1– 522^, JIP3^FL^, JIP3^1–582^, JIP3^1–560^ and JIP3^1–185^. **(C)** Quantification of the number of processive events/μm microtubule/minute with the mean ± S.D. plotted. The total number of events analysed were 1364 (HOOK3^1–522^), 99 (JIP3^FL^), 348 (JIP3^1–582^), 537 (JIP3^1–560^) and 513 (JIP3^1–185^). **(D)** An AlphaFold2 prediction of two copies of JIP3 (residues 1-600) showing an interaction between JIP3^helix^ (residues 570-582) and the RH1 domain. The JIP3^helix^ contains the same FFxxL motif as the DLIC^helix^ (residues 427-439 of the DLIC2 isoform) (bottom left), which has been shown to bind the RH1 domain^37^ (bottom right). **(E)** Coomassie Blue-stained SDS-PAGE gel of purified recombinant protein mixtures prior to the addition of glutathione agarose resin (left) and of proteins eluted from glutathione agarose resin after GST pull-down (right). The RH1 domain construct was composed of JIP3^1–108^ and JIP3^helix^ construct was composed of GST-JIP3^563–586^. The asterisks denote the bands corresponding to degradation products. **(F)** Kymograph of TMR-dynein-dynactin-LIS1 in the presence of JIP3^1– 582^ mut (left). Quantification of the number of processive events/μm microtubule/minute with the mean ± S.D. plotted. The total number of events analysed were 568. Statistical significance in comparison with JIP3^1–582^ and JIP3^1–560^ is depicted. The *in vitro* motility assays were performed with three technical replicates and statistical significance was determined using ANOVA with Tukey’s multiple comparison.

One putative family of adaptors that does not appear to conform with these rules includes JIP3 (c-Jun N-terminal kinase-interacting protein 3)^1^. This is an important protein for trafficking neuronal lysosomes and autophagosomes^30–33^ and its defects can lead to lysosome accumulations, increased levels of amyloidogenic amyloid precursor protein and neurological diseases^32,34–36^. JIP3 contains an N-terminal RILP Homology 1 (RH1) domain which binds DLIC^helix^^37^. However, its N-terminal coiled coil, which is called Leucine Zipper 1 (LZ1)^30^, is roughly half the length of other known dynein adaptors (Figure 1A). This, together with the absence of a defined HBS1 or Spindly motif, made it initially unclear whether JIP3 can directly activate dynein^1^ and, if so, what might be the underlying mechanism.

Along with the adaptor, DDA complex formation also requires interactions between dynein and dynactin. Dynactin is built of a short actin-related filament capped by a pointed end complex, and contains a shoulder domain from which the long p150^Glued^ (p150) projection or arm extends^28,38–40^. Cryo-electron microscopy (cryo-EM) showed how grooves on dynactin’s filament bind the N-terminal DHC tails of dynein dimers. This stabilises dynein’s C-terminal motor domains in an open parallel conformation and contributes to their activation^14^. In addition, the DIC N-terminus (DIC-N) interacts with p150^41–46^, which is critical for dynein functions in cells^47^ and the formation of DDA complexes *in vitro*^48^. However, the p150 adopts an ‘autoinhibited’ folded-back conformation^38,40,49^ which is thought to block DIC-N binding. This leads to questions of when DIC-N binds p150 and why this interaction is critical for dynein’s activity.

LIS1 was first identified by mutations which cause Lissencephaly, a brain developmental disorder caused by defects in neuronal migration^50^. It was shown to be important for dynein function in a wide range of eukaryotes^51–58^ and recent evidence suggests it stimulates DDA complex formation^59–63^. The current model is that LIS1 achieves this by binding dynein’s motor domains^64–67^ and opening up an autoinhibited form of dynein referred to as the “phi” particle^60,62,63,68^. However, LIS1 can still stimulate DDA complex formation even in a dynein mutant where the phi-particle is constitutively open^59,60^. Furthermore, the phi-opening model does not explain other reported LIS1 functions, such as aiding recruitment of dynein-dynactin to microtubule plus-ends^56,61,69^. In the absence of structural information of LIS1 in the context of the whole DDA complex, it is therefore not clear if the phi-opening model is sufficient to explain its different roles.

In this study, we initially set out to investigate whether JIP3 can activate dynein-dynactin. This led us to identify a mechanism by which full-length JIP3 is autoinhibited and find the minimal regions involved in dynein activation. We used cryo-EM to determine the structure of a complex containing dynein, dynactin, JIP3 and LIS1 bound to microtubules. This revealed the motifs JIP3 uses to recruit dynein and explains why JIP3’s short coiled coil is sufficient for activation. It also allowed us to resolve critical regions of dynein-dynactin that were previously elusive. The dynein light chains (LC8 and TCTEX1) are tucked underneath the dynein heavy chain positioning the DIC-N near the dynein motor. Furthermore, LIS1 stabilises much of the p150 arm allowing us to see its interaction with DIC-N and understand how it stimulates DDA assembly. We also discovered that LIS1 directly tethers p150 to the dynein motor domain and show that this is essential for dynein activity *in vitro* and in cells. Together, our study provides the molecular mechanism for DDA complex assembly and how LIS1 stimulates this process.

## Results

### JIP3 is an autoinhibited dynein activating adaptor

To test if JIP3 can directly activate dynein-dynactin, we quantified its ability to stimulated processive movement using *in vitro* motility assays^13^. We included LIS1 in all experiments to increase the robustness of dynein activation^59,60^. Under our assay conditions, the known activating adaptor construct^12,70^ HOOK3^1–522^ showed 1.07 ± 0.3 (SD) processive events/μm/min (Figure 1B, C, Table S1). In contrast, full-length JIP3 (JIP3^FL^) displayed only a small number of processive events (Figure 1B, C) which were not significantly higher than in the absence of any adapter (Figure S1A, Table S1). This is reminiscent of other activating adaptors which are autoinhibited in their full-length form and can be activated by truncation of their C-terminal segments^26,71^. We therefore generated JIP3 truncations and tested their ability to activate dynein. JIP3^FL^ contains an N-terminal RH1 domain, three coiled coils, (LZI, LZII and RH2) followed by a WD40 domain (Figure 1A). Removing the WD40 domain (JIP3^1– 582^) led to an increase in the number of processive events, although this was not statistically significant (Figure 1B, C). In contrast, shorter JIP3 constructs, JIP3^1–560^ and JIP3^1–185^, resulted in many long-distance dynein runs (Figure 1B). The frequency of processive events (Figure 1C, S1B), as well as the velocity and run lengths (Figure S1C, D), were comparable to HOOK3^1–522^. This demonstrates that the level of activation by JIP3 and the motile properties of resulting DDA complexes are similar to well-studied dynein adaptors like HOOK3.

The significant increase in dynein activation upon deletion of residues 561-582 suggests this region plays a role in autoinhibiting JIP3. An AlphaFold2 prediction of JIP3 (residues 1-600) shows a high confidence interaction between a helix within the 561-582 region (JIP3^helix^) and the RH1 domain (Figure 1D, S1E). This resembles the previously reported^37^ interaction of the RH1 domain with the DLIC^helix^ (Figure 1D, S1E). To test the AlphaFold2 model, we performed pull-downs with purified proteins. Our data demonstrate that a JIP3^helix^ construct (GST-JIP3^563–585^) directly binds the RH1 domain (JIP3^1–108^) (Figure 1E, S1F). This interaction was abrogated by an RH1-domain mutant (V60Q)^37^ which is unable to bind DLIC^helix^ (Figure 1E, S1F). Furthermore, the JIP3^helix^ contains the same FFxxL motif as the DLIC^helix^ (Figure 1D) and mutating residues in this motif (JIP3^F576A^ or JIP3^L579A^) breaks the JIP3^helix^-RH1 interaction (Figure S1F). Together these results show that JIP3^helix^ binds to the RH1 domain in a similar manner to DLIC^helix^.

We next tested the effect of disrupting the JIP3^helix^-RH1 interaction on the ability of JIP3^1–582^ to activate dynein. Mutating the two phenylalanine residues to alanine in the JIP3 FFxxL motif (JIP3^1–582^ mut) resulted in 1.15 ± 0.23 (SD) events/μm/min, which is comparable to that observed for JIP3^1–560^ and JIP3^1–185^ (Figure 1F). Collectively these data show that JIP3 is autoinhibited by an intramolecular interaction between the JIP3^helix^ and the RH1 domain that likely hinders dynein’s DLIC^helix^ from binding.

### Structure of dynein-dynactin bound to JIP3 and LIS1

To understand how JIP3 recruits dynein-dynactin, we determined the structure of these complexes using cryo-EM. We prepared complexes of dynein-dynactin with JIP3^1–185^ (DDJ^1–185^) or JIP3^1– 560^ (DDJ^1–560^) and decorated them on microtubules in the presence of a non-hydrolysable nucleotide analogue, AMPPNP. As in our *in vitro* motility assays, we prepared these DDJ complexes in the presence of LIS1 to stimulate complex formation.

For structure determination, we used a cryo-EM processing pipeline involving microtubule signal subtraction^72^ followed by focused 3D classification and 3D refinement^27^. The cryo-EM datasets from the two different JIP3 constructs were later combined to better resolve regions of the complex present in both DDJ^1–185^ and DDJ^1–560^ structures (Figure S2).

Our composite structure, generated from locally refined regions of the complex, shows that a single JIP3 adaptor recruits two dynein dimers per dynactin (Figure 2A, S2). As in previous structures^14,27,40,70^, the dyneins bind dynactin via their tail regions. When viewed with dynein walking towards the reader, dynein-A with its heavy chains A1 and A2 are located on the left whereas dynein-B is located on the right (Figure 2B). The dynein-B motor domains are both in contact with the microtubule via their stalks, whereas the dynein-A motors are lifted away from the microtubule surface.

**Figure 2.**
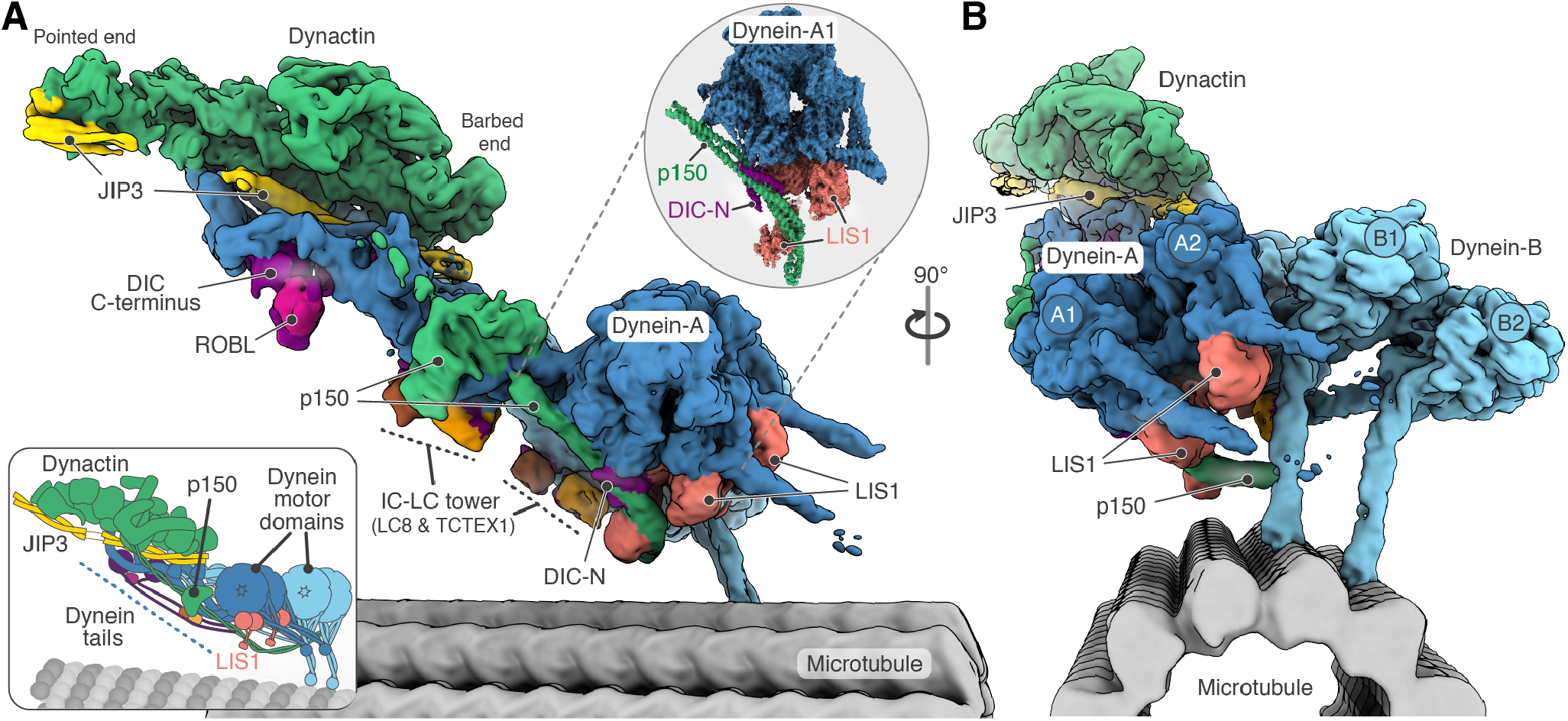
Cryo-EM structure of dynein-dynactin-JIP3-LIS1 on microtubules. **(A), (B)** Composite density map of dynein-dynactin bound to JIP3 and LIS1 overlaid on 13 protofilament microtubule. The zoom-in (grey circle) shows the locally refined map of dynein-A1 bound to p150, DIC-N and LIS1. Schematic representation of the complex is shown on the bottom left.

The structures contained ordered density for many functionally important parts of dynein-dynactin that were too flexible to visualize in previous DDA complex structures. These include much of dynactin’s p150 arm, structured segments of the DIC bound to the dynein light chains LC8 and TCTEX1 (together referred to as the IC-LC tower) and the DIC-N. Furthermore, we found two LIS1 dimers stably bound to the complex, one on each motor domain of dynein-A. To our surprise, one of these LIS1 dimers also contacts p150, thereby directly linking dynein to dynactin.

In the following sections we first explain how JIP3 recruits dynein-dynactin. We then focus on the unanticipated interactions between dynein, dynactin and LIS1 and their role in DDA complex formation.

### Molecular basis of dynein-dynactin recruitment by JIP3

In our structure, JIP3 residues 24-185, containing the RH1 domain and an ∼18 nm long LZI coiled coil, bind along the cleft between the dynein tails and dynactin (Figure 3A, B). The C-terminal JIP3 residues 374-550, encompassing the LZII and RH2 coiled coils, bind at dynactin’s pointed end. The intervening ∼200 amino acids are disordered and therefore not visible. This arrangement of JIP3 differs from other structurally-characterized dynein adaptors, whose long-coiled coils bind along the length of the dynactin filament extending all the way to the pointed end^27,28,70^.

**Figure 3.**
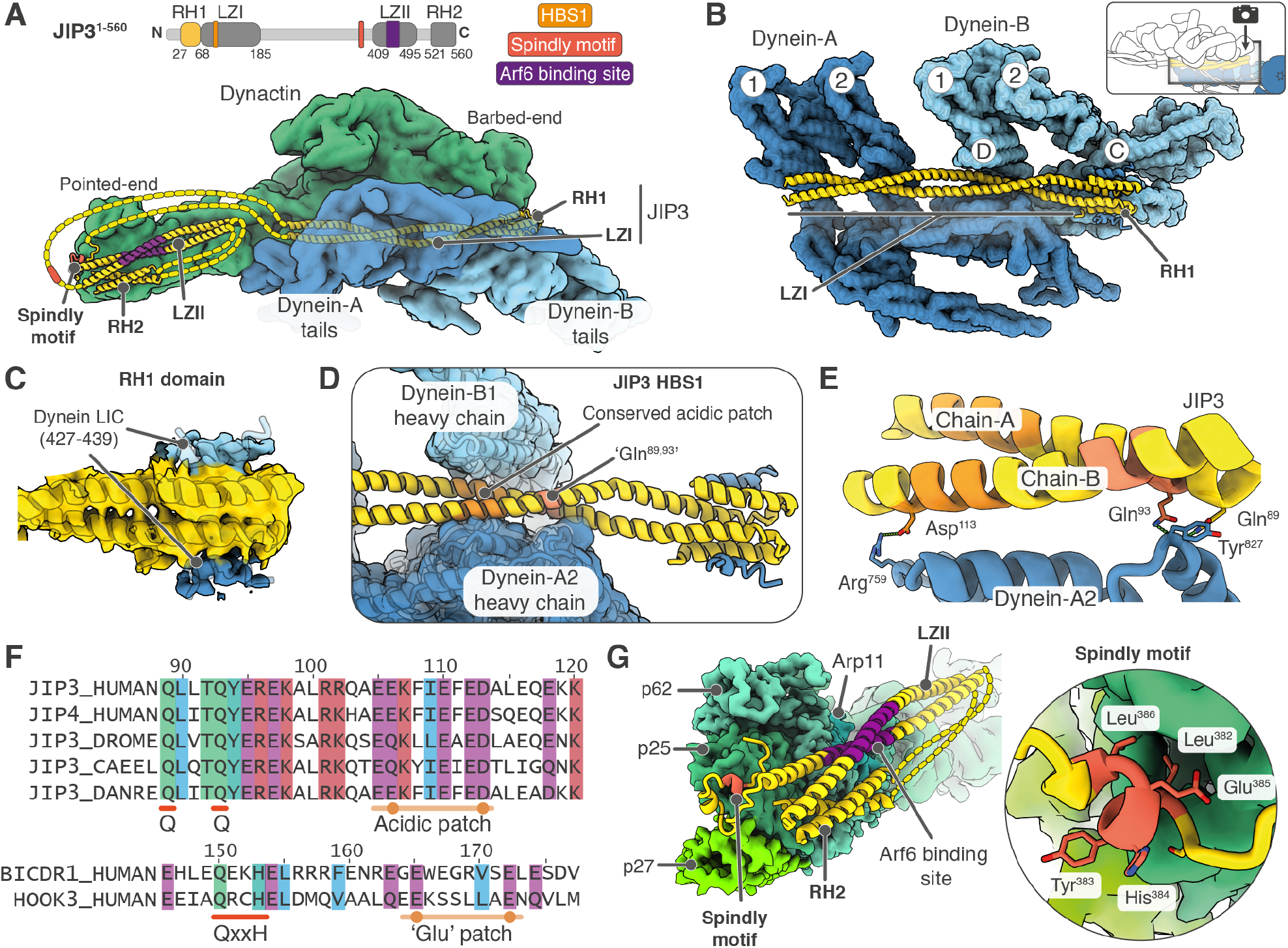
JIP3 interactions with dynein-dynactin. **(A)** Overall organisation of JIP3 bound to dynein-dynactin. The unresolved regions which are predominantly unstructured are depicted as a dotted line. **(B)** Trajectory of JIP3 along the dynein heavy chain tails is shown using a top view of the complex. Dynactin segments are removed to aid visualization. **(C)** The N-terminal RH1 domain (yellow) has extra density on either side which is explained by the DLIC C-terminal helix (blue). The AlphaFold2 prediction of this interaction is displayed as cartoon inside the density. **(D)** JIP3 is sandwiched between heavy chains of dynein-B1 and dynein-A2. JIP3 contains an HBS1 which interacts with the dynein heavy chains using a glutamate residue followed by an acidic patch. **(E)** JIP3 interaction with the dynein-A2 is shown. **(F)** Multiple sequence alignment of the JIP3 HBS1 from different organisms is shown. The residues involved in interactions with dynein are highly conserved. Sequences of HBS1 from BICDR1 and HOOK3 are shown for comparison. **(G)** JIP3 segments bound at the dynactin pointed end are shown. The JIP3 Spindly motif binds to the p25 subunit of dynactin whereas the LZII and RH2 foldback on each other and are bound along the p25 and p62 subunits (left). Spindly motif residues bound at the p25 subunit are shown on the right.

The interactions made by the RH1 domain and LZI are sufficient to promote dynein motility since the JIP3^1–185^ construct only contains those regions and is able to activate dynein in motility assays (Figure 1B). The RH1 domain docks above the dynein-B1 tail (Figure 3B) and shows extra density consistent with two copies of DLIC^helix^, one on each side^37^ (Figure 3C). Distance constraints make it most likely that one DLIC^helix^ comes from dynein-A and the other from dynein-B. The LZI coiled coil is sandwiched between the dynein-A2 and B1 tails (Figure 3B, D). Within the LZI, residues Gln^89^ and Gln^93^ interact with dynein-A2 via residue Tyr^827^, and a downstream patch of acidic residues interacts with both dynein-A2 (residues 420-460) and dynein-B1 (residues 759- 830) (Figure 3D, E). These JIP3-dynein interactions constitute its HBS1 and are analogous to the HBS1 motif (a QxxY/H followed by a patch of glutamates) found in BICDR1 and HOOK adaptor families^25,27^ (Figure 3F). However, the Gln^93^ of JIP3 sits on the opposite side of dynein Tyr^827^ compared to the equivalent Gln^150^ in BICDR1 (Figure 3E and S3A). Furthermore, the downstream acidic patch is spaced differently with respect to the glutamine residue in these two adaptors (Figure 3F). These differences in HBS1- dynein interactions are due to the different rotational orientation of the JIP3 coiled coil compared to BICDR1 (Figure S3A), and illustrate the variability in how adaptors can recognise the DHCs.

In the longer JIP3 construct (JIP3^1–560^) we observe additional density corresponding to two coiled coils (∼12 nm and ∼6 nm in length) docked against dynactin’s pointed end complex subunits Arp11, p25, p27 and p62 (Figure S3B). Based on the dimensions, we assigned the longer coiled coil to the LZII and the shorter one to the RH2 domain (Figure 3G). Connected to LZII is a stretch of density containing a short alpha helix which docks onto the p25 subunit of dynactin (Figure 3G, S3B). This is located at the site on p25 where the Spindly motif (LΦXEΦ, where Φ is hydrophobic) binds in other adaptor structures^27,28^. Upon searching the sequences flanking LZII, we found JIP3 contains a sequence, LYHEL (residues 382-386), that matches the consensus of the Spindly motif and explains our cryo-EM density (Figure S3B). To support this assignment, we performed gel filtration assays to assess the interaction of JIP3 with a recombinantly-expressed pointed end complex^29^. These showed that whereas JIP3^1–560^ binds the pointed end, mutations in the putative Spindly motif (L382A, Y383A and E385A) disrupt this interaction (Figure S3C). Together, our data shows that JIP3 contains a previously unidentified Spindly motif that is critical for the JIP3-pointed end interaction.

LZII is important for JIP3 function due to its interaction with the small G-protein Arf6, thereby connecting the adaptor to membrane cargos^30^. However, the crystal structure of Arf6 bound to the JIP3 paralog, JIP4^73^, suggests Arf6 would sterically hinder LZII’s interaction with the pointed end (Figure S3D). This implies that either 1) Arf6 prevents JIP3 binding to dynactin or 2) Arf6 causes LZII to detach from the pointed end, but JIP3 can remain attached via other means. To test these scenarios, we performed pull-downs using a JIP3 fragment (mouse JIP3^185–505^) containing the Spindly motif and LZII (Figure S3E). We found this fragment can simultaneously bind both Arf6 and the recombinantly-expressed pointed end complex. These interactions are specific and mutually independent. A mutation in LZII (L439P, corresponding to a human disease mutation L444P^36^) disrupts Arf6 binding without affecting the pointed end interaction. Conversely, mutating the Spindly motif (L383A/E386A) disrupts the pointed end interaction without affecting Arf6 binding (Figure S3E). Together, this suggests that when JIP3 is on cargos, it binds the pointed end complex via its Spindly motif whereas the LZII binds Arf6 and is detached from the dynactin (Figure S3F).

### Dynactin’s p150 arm binds DIC-N, DHC and LIS1

The dynein-dynactin machinery contains two long flexible regions, the p150 arm and the DIC N-terminus, both of which are important for complex assembly and function^41,42,48,74–76^ (Figure 4A). The p150 arm extends as a coiled coil (CC2) from the dynactin shoulder^28,38–40^. It continues from C-to N-terminus as a globular intercoiled domain (ICD), followed by two coiled coils (CC1B and CC1A), a basic-rich region and CAP-Gly domains, the latter two mediating p150’s interaction with microtubules^28,38–40^. CC1B is the binding site for the very N-terminal ‘region 1’ helix of DIC-N^44,45^, although the exact interaction site is not known. Previous work showed CC1A and CC1B fold back on each other in isolated dynactin^38,40,49^. This led to the idea that the folded-back CC1A/B hairpin is in an autoinhibited conformation that is opened upon DIC-N binding, although what role this plays in DDA assembly is also not clear.

**Figure 4.**
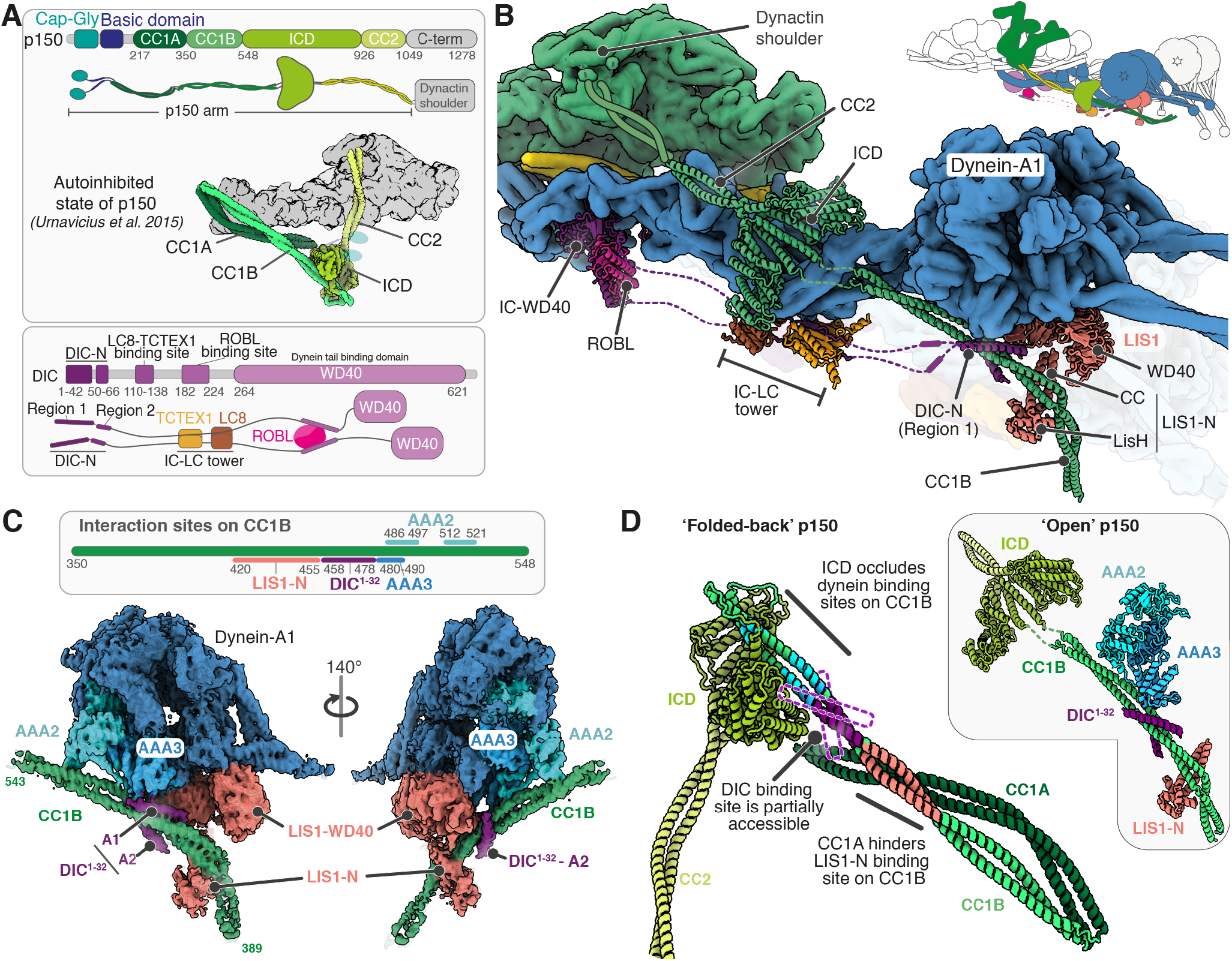
Organisation of p150 and dynein intermediate chain. **(A)** Domain architecture of p150 (top) and its organisation in the autoinhibited conformation is shown. Domain architecture of dynein intermediate chain and the interaction sites for dynein light chains are depicted (bottom). **(B)** Cartoon representation of p150 (green), dynein intermediate chain (magenta), dynein light chains (ROBL (pink), LC8 (brown) and TCTEX1 (golden)) and LIS1 (coral) bound to dynein-A1 (blue; depicted as surface). Unresolved segments are depicted as dotted or solid lines. **(C)** Schematic of interaction sites on p150-CC1B (top) and cryo-EM density of dynein-A1 motor domain bound by CC1B and LIS1 (bottom). CC1B is further bound by DIC-N region 1 (residues 1-32). **(D)** The binding sites for dynein motor domain, DIC^1–32^ and LIS1-N on CC1B are mapped on the model of the autoinhibited p150. The open state of p150 solved in this study is shown on the right for comparison. The area covered by the two DIC^1–32^ helices when bound to CC1B is shown using dotted lines.

In our JIP3- and LIS1-bound dynein-dynactin complex, p150 stably docks onto dynein allowing us to see much of its length. The ICD binds to the dynein-A1 heavy chain (residues 1160-1400) (Figure 4B, S4A). The CC1A/B hairpin has opened up, consistent with autoinhibition being relieved. CC1B itself contacts the dynein-A1 motor (Figure 4B). The N-terminal segments of p150 including CC1A, basic-rich region and CAP-Gly domains are not visible in our structure, however the interaction of CC1B with the dynein would position them close to the microtubule.

Our structure shows that CC1B acts as a major interaction hub (Figure 4C). Its C-terminal end (residues 480-521) contacts the dynein-A1 motor. Dynein motor domains are built of a ring of six AAA domains (AAA1 – AAA6) and here we see CC1B specifically contacts AAA2 and AAA3. The adjacent, N-terminal section of CC1B (residues 458-478) binds two alpha helices, one on each side of its coiled coil. We assign these helices to DIC-N based on previous reports^41,44–46^. Their length, together with an AlphaFold2 prediction of DIC-N bound to CC1B suggests they correspond to the first part of DIC-N Region1 (residues 1-32) (Figure 4C, S4B).

Immediately N-terminal to the DIC-N binding site on CC1B is a section (residues 420-455) which binds a LIS1 molecule (Figure 4B, C). LIS1 contains an N-terminal LisH domain and coiled-coil region (together referred to as LIS1-N) required for its dimerization^77^ and a C-terminal WD40 domain which binds the dynein motor^67,78,79^. Of the two LIS1 dimers in our structure, the one which binds dynein-A1, contacts CC1B via its LIS1-N domain. Notably, the CC1B coiled coil curves around LIS1-N and is redirected to run under the dynein motor domains (Figure 4B, C). The interaction sites on CC1B/DIC-N/LIS1-N are highly conserved (Figure S4C) and are also supported by an AlphaFold2 prediction of these segments (Figure S4B).

### DIC binding initiates opening up of dynactin’s p150 arm

The structure raises the question of which interactions are required to open up the CC1A/B hairpin. To address this, we used AlphaFold2 together with previously published cryo-EM maps and crosslinking data^28,40^ to generate a model of the autoinhibited form of p150 (Figure 4D, S4D). We find that CC1B is docked onto the ICD in this model, but detached and connected only by a flexible linker in the open p150 (Figure 4D). When docked, the ICD blocks the dynein motor binding regions on CC1B, showing an unanticipated role of the ICD in p150 autoinhibition. The model further shows that the previously identified hairpin, formed by CC1A binding to CC1B, is incompatible with LIS1-N binding. In contrast, the DIC-N binding site is partially accessible. Comparison of the two p150 states suggests that DIC-N binding would destabilise the autoinhibited p150 as one DIC-N copy would partially clash with the ICD whereas the other would displace CC1A (Figure 4D). This implies that DIC-N binding must occur first to overcome the autoinhibition and allow CC1B to bind LIS1 and the dynein motor.

In support of the above model we find LIS1 or LIS1-N are unable to pull-down with a CC1A/B construct (GST-CC1A/B), whereas they do bind GST-CC1B (Figure S4E, F). In contrast, a monomeric DIC-N construct (MBP-DIC-N) can bind GST-CC1A/B, consistent with the partial accessibility of its binding site. However, the binding of MBP-DIC-N to either GST-CC1A/B or GST-CC1B was sub-stoichiometric. We hypothesised that this was due to the lower affinity of monomeric DIC-N compared to a bivalent form found in dynein^41,80^. To address this, we changed the assay so that the beads are decorated with MBP-DIC-N instead. This has the effect of increasing the local concentration and under these conditions we pulled down CC1A/B in stoichiometric amounts. When DIC-N-decorated beads are incubated with CC1A/B and LIS1 or LIS1-N, we found a complex formed between all three components, providing evidence that DIC-N binding is required for LIS1 to interact with p150 (Figure S4G). Together, our structure and pull-down assays suggest DIC-N binds p150 first and that the function of this interaction is to open up CC1A/B hairpin allowing it to bind the dynein motor and LIS1.

### The IC-LC tower promotes the CC1B/DIC-N interaction

Our cryo-EM structure showed two distinct globular densities below both dynein-A and B heavy chains (residues 1160-1230) (Figure S5A). The resolution of our maps in this region was ∼9 Å which is sufficient to identify them as belonging to the IC-LC tower based on its crystal structure^80,81^ (Figure S5B). It is composed of DIC residues 110-140 and two copies each of LC8 and TCTEX1. We find that for each IC-LC tower, LC8 binds to the dynein heavy chain-1 (A1/B1) while TCTEX1 binds to the dynein heavy chain-2 (A2/B2) (Figure S5A). Such a configuration suggests that the IC-LC tower plays a structural role. Firstly, the interaction with the DHCs reinforces the dynein tails parallel to one another in each dynein dimer. Secondly, the docking of the IC-LC tower positions DIC-N so it tethers the p150 close to the dynein-A motor domain.

### LIS1 stabilizes the pre-powerstroke state of dynein-A

In our structure, the two dynein dimers adopt different conformations (Figure 5A). The LIS1- bound dynein-A motors have a bent linker docked onto their AAA2/AAA3 domains, indicating they are in pre-powerstroke state (low microtubule affinity) (Figure 5B). On the other hand, the dynein-B motor domains have a straight linker docked onto AAA5 indicating they are in the post-powerstroke state (high microtubule affinity) (Figure 5B). In dynein-A motors we find the nucleotide density in both the AAA1 and AAA3 pockets is consistent with ADP (Figure 5B, S5C). In contrast, the dynein-B motors contain ADP and AMPPNP in AAA1 and AAA3, respectively (Figure 5B, S5D). The nucleotide state of dynein-A is surprising since the presence of ADP in AAA1 typically corresponds to a high microtubule affinity state^82,83^, whereas dynein-A motors are detached from the microtubule in our structure. Our data suggest that LIS1 overrides the dynein conformation dictated by the bound nucleotides, stabilizing the pre-power stroke state. The result of stabilizing this conformation is to allow p150 to dock onto the dynein heavy chain as described above. The regions of dynein heavy chain that bind to p150 are only accessible when the linker is bent and occluded when it is straight (Figure 5C). Thus, LIS1 aids p150 docking via its ability to drive dynein-A into a pre-powerstroke conformation.

**Figure 5.**
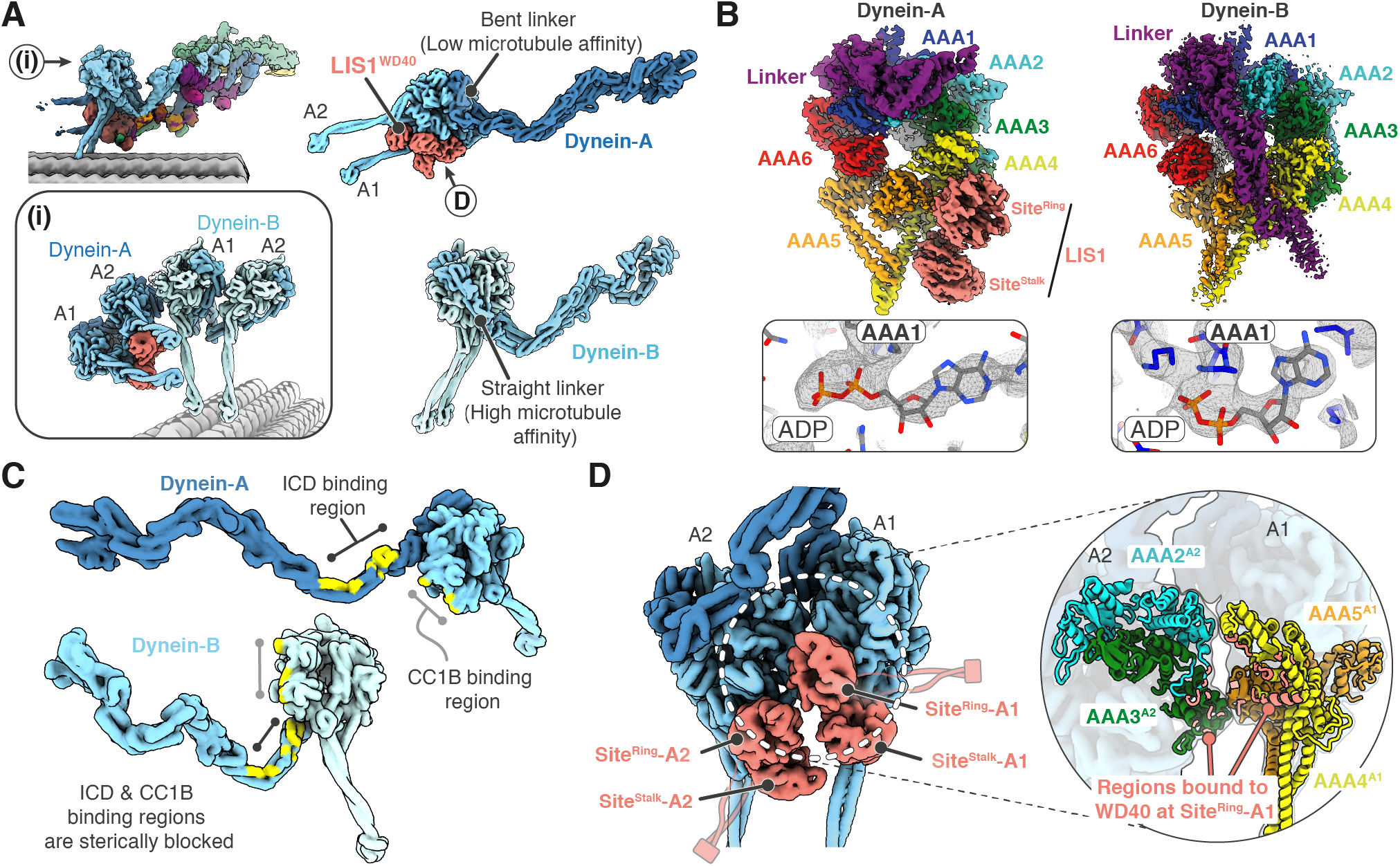
LIS1 stabilizes the pre-powerstroke state of dynein-A. **(A) (i)** Surface representation of LIS1 bound dynein-A and microtubule bound dynein-B. Dynein conformation and linker position is shown on the right. **(B)** Cryo-EM density of dynein-A and -B motor domains where the different subdomains and bound proteins are highlighted. **(C)** Regions involved in interacting with the ICD and CC1B segments of p150 are mapped onto the heavy chain of dynein-A (top) and dynein-B (bottom). **(D)** Surface representation of dynein-A motor domains with bound LIS1-WD40 domains (left). Interaction between dynein-A1 and A2 motor domains is shown (right) and the residues at the A1-A2 interface interacting with LIS1 are mapped (coloured in coral).

In our structure, the two LIS1 dimers contact the dynein-A motors via their WD40 domains. They bind at two sites: one at the interface between AAA3-AAA4 (Site^ring^) and the other at the base of the stalk (Site^stalk^) (Figure 5B). These are consistent with previous structures of LIS1 bound to isolated motor domains^64–67^. In addition, LIS1 bound in the context of the full dynein dimer reveals an additional, previously unanticipated interaction. A molecule of LIS1 sits in the cleft between the A1 and A2 motor domains, with the Site^ring^ WD40 domain on dynein-A1 contacting the AAA3 domain of dynein-A2 (Figure 5D). This stabilizes dynein-A2 docking on AAA5 of dynein-A1 (Figure 5D) and keeps the two motor domains parallel. Thus, LIS1 can affect both the conformation and organisation of the two dynein motors.

### LIS1 – p150 interaction is important for complex assembly

The ability of LIS1 to stimulate DDA complex formation is attributed to its WD40 domains binding the dynein motor and disrupting its autoinhibited phi particle^62,63,68^. However, the interaction between LIS1-N and p150-CC1B identified in this study raises the possibility that LIS1 has additional roles in dynein activation. To understand if the LIS1-N/CC1B interaction is important, we first tested if the LIS1-WD40 domains alone are sufficient to activate dynein transport in cells. For this, we used a mitochondria relocation assay where we generated a Flp-in T-REX HeLa cell line expressing a GFP-labelled N-terminus of the BICD2 adaptor (BICD2N) fused to a mitochondria targeting sequence (MTS) (Figure 6A). In this cell line, mitochondria are clustered to the perinuclear region in a BICD2 dependent manner, and knockdown of LIS1 or dynein increases their spread (Figure S6A, B). Importantly the LIS1 knockdown can be rescued by electroporating the cells with LIS1 protein (Figure 6B, C). The LIS1 used in this assay had a C-terminal SNAP tag (LIS1-SNAP) for detection, but behaved identically to wild-type LIS1 (Figure S6C).

**Figure 6.**
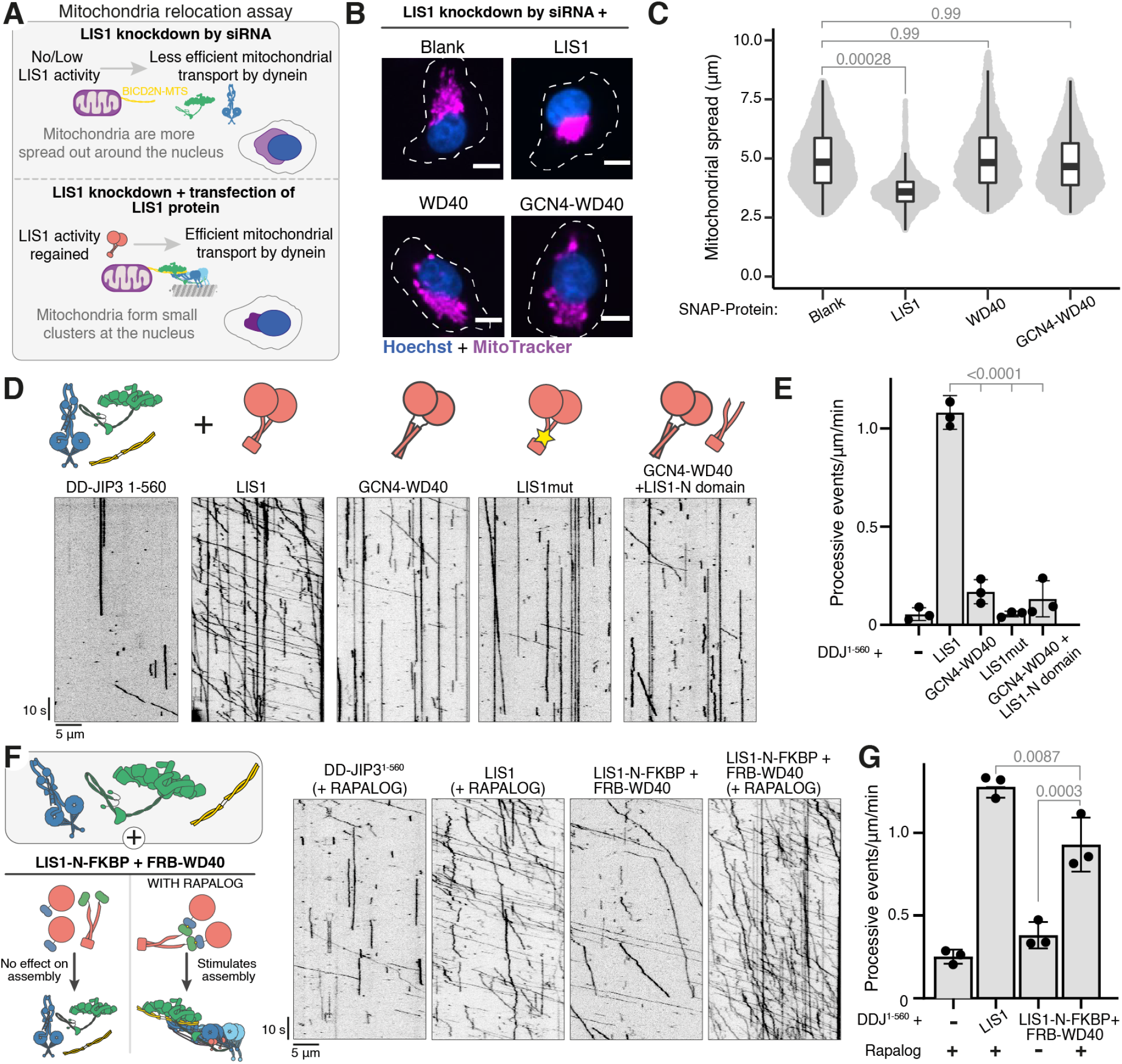
LIS1-N binding to p150 is important for dynein activation. **(A)** A schematic of the mitochondrial relocation assay is shown (left). **(B)** Representative images showing the distribution of mitochondria (magenta) in LIS1 siRNA-treated HeLa GFP-BICD2N-MTS cells electroporated with different SNAP-tagged LIS1 proteins. Scale bars represent 10 μm. **(C)** Quantification of mitochondrial spread (μm) in LIS1 knockdown HeLa GFP-BICD2N-MTS cells electroporated with different LIS1 proteins. Data are plotted from 3 biological replicates. The total cells quantified were 3922 (blank), 3830 (LIS1-SNAP), 2541 (WD40-SNAP) and 3655 (GCN4-WD40-SNAP). **(D)** Kymographs of TMR-dynein-dynactin-JIP3^1–560^ in the presence of (from left) blank, LIS1, GCN4-WD40, LIS1^mut^ and GCN4-WD40+LIS1-N. Cartoons depicting the LIS1 construct used are shown above each kymograph. **(E)** Quantification of the number of processive events per μm microtubule per minute with the mean ± S.D. plotted. The total number of movements analyzed were 74 (blank), 1592 (LIS1), 352 (GCN4-WD40), 116 (LIS1^mut^), and 225 (GCN4-WD40 + LIS1-N). Data are plotted from 3 technical replicates. **(F)** Schematic summarizing the effect on dynein assembly when LIS1-N and WD40 domains are present on the same molecule (left). Kymographs of TMR-dynein-dynactin-JIP3^1–560^ in the (from left) absence of LIS1, with LIS1, with FRB-WD40+LIS1-N-FKBP without and with rapalog are depicted (right). **(G)** Quantification of the number of processive events per μm microtubule per minute with the mean ± S.D. plotted. The total number of movements analysed were 283 (blank), 675 (LIS1), 388 (FRB-WD40+LIS1-N-FKBP) and 665 (FRB-WD40+LIS1-N-FKBP with rapalog). Data are plotted from 3 technical replicates. Statistical significance was determined using ANOVA with Tukey’s multiple comparison.

To test whether the WD40 domain alone can rescue LIS1 knockdown we used two constructs: one with a monomeric WD40 (WD40-SNAP) and a second dimeric construct where the LIS1-N dimerization domain was replaced by a GCN4 coiled coil (GCN4-WD40-SNAP) (Figure S6D). Both constructs can bind to the dynein motor domain (Figure S6E) but lack the ability to interact with p150-CC1B (Figure S6F). The constructs were electroporated into LIS1 knockdown cells (Figure S6G), but in contrast to wild-type LIS1, neither were able to rescue perinuclear mitochondrial clustering (Figure 6B, C). This suggests the LIS1-N plays a critical role in dynein activation.

To dissect the role of the LIS1-N/CC1B interaction in LIS1 function, we used our *in vitro* motility assay to check for formation of active DDJ^1–560^ complexes. Consistent with our cellular observations, the presence of wild-type LIS1 led to multiple processive dynein runs whereas the monomeric WD40 or GCN4-WD40 constructs were unable to activate dynein under the conditions used in this assay (Figure 6D, E, S6H).

Additionally, we tested a construct in which the CC1B interacting interface of LIS1 is disrupted by mutation of 13 residues in the LIS1-N region (Figure S6F, I). This construct (LIS1^mut^) is a dimer (Figure S6D) and binds the dynein motor (Figure S6E) but was unable to activate dynein transport (Figure 6D, E), showing that CC1B binding by LIS1 is important for LIS1 activity.

We found that LIS1-N alone was also unable to stimulate dynein activation, consistent with previous observations showing that the WD40- dynein interaction is necessary^59^ (Figure S6H). Furthermore, adding both LIS1-N and GCN4- WD40 constructs together did not increase in the number of processive dynein runs (Figure 6D, E). This indicates that the LIS1-N and WD40 domains must be linked together in order to activate dynein.

To directly test if LIS1 activity requires a connection between LIS1-N and WD40, we used the rapalog inducible FKBP-FRB system. LIS1-N was fused to FKBP and the WD40 domain to FRB. In the absence of rapalog, we observed similar numbers of processive dynein events as in the absence of LIS1 (Figure 6F, G). Upon addition of rapalog, we observed a significant increase (∼3-fold) in dynein activation which resulted in ∼73% as many processive events as observed for wild-type LIS1 (Figure 6F, G). Together with our structure, this finding suggests that dynein activation requires LIS1 to simultaneously bind the dynein motor domain and the dynactin p150.

## Discussion

### Activation of the dynein machinery by JIP3

We have shown that JIP3 can activate long-range dynein movement despite having a much shorter coiled coil than other dynein cargo adaptors^1^. A minimal fragment (JIP3^1–185^) containing just the RH1 domain and LZI short coiled coil is sufficient for activation (Figure 1). The LZI coiled coil spans the minimum distance required to interact with the tails of two dyneins (Figure 3B) providing an explanation for its length within JIP3. It also suggests that an interaction with the dynactin pointed end complex is not required for dynein activation. In fact, there appear to be few, if any, interactions between JIP3^1–185^ and the dynactin complex. This suggests that the role of adaptors is to orient and stabilize the binding of two dyneins to each other so that their heavy chain tails can interact with the grooves in dynactin’s filament. This part of our work was previously released in our preprint^84^ and is consistent with other contemporary studies reporting that JIP3 can activate dynein-dynactin^85,86^.

Our study also shows full-length JIP3 is autoinhibited (Figure 1). This is due to a short helix (JIP3^helix^: residues 570-582), just C-terminal to the RH2 domain, which likely blocks dynein’s DLIC^helix^ from binding to JIP3’s RH1 domain. This mechanism is distinct from that reported for other coiled-coil adaptors. For example, in the case of BICD2^87–89^ and Spindly^26^ inhibition is due to a C-terminal coiled coil directly binding the N-terminal coiled coil. Interestingly, a recent study reported dynein-dynactin movement with full-length JIP3^85^. However, this work used cell lysates which likely contain proteins that recruit JIP3 to cargos such as Arf6 (binds to LZII)^30^ or Rab36/Rab10 (bind to RH2 domain)^90,91^. These factors likely disfavour JIP3’s autoinhibited conformation and promote dynein activation.

### Role of dynactin pointed end interactions

Our cryo-EM structure of dynein-dynactin bound to the longer JIP3^1–560^ construct allowed us to identify JIP3’s Spindly motif (Figure 3). The disordered region between the dynein-binding coiled coil (LZI) and the Spindly motif is much longer in JIP3 than other adaptors. Furthermore, there is a gap of 24 residues between the Spindly motif and the next closest alpha helical region (LZII) in JIP3, whereas in other adaptors, it is situated right next to a long alpha helix. These differences explain why JIP3’s Spindly motif was not identified previously.

The JIP3^1–560^ cryo-EM structure shows the LZII and RH2 coiled coils bind dynactin’s pointed end (Figure 3). Our data suggest, however, that Arf6 binding to LZII is incompatible with this docking. Thus, when JIP3 binds cargo it is likely to only use its Spindly motif to contact the pointed end. We anticipate, similarly, that the RH2 docking is also disrupted when it is bound to Rab10^91^ or Rab36^90^. This agrees with previous data showing that RH2 is not required for JIP3 to bind dynactin^30^.

When LZII is detached, we calculate that the distance from the Spindly motif to the Arf6 binding site is around 7.5 nm (Figure S7A). This compares with a distance of ∼46 nm between the Spindly motif and Rab6 binding site on the adaptor BICD2. Other adaptors show equivalent distances which lie between these two values. This means the dynein-dynactin-JIP3 complexes are positioned much closer to cargo membranes than some other DDA complexes. For organelles that could simultaneously recruit multiple types of adaptors this would allow more motors to attach the vesicle to the microtubule than would be possible if all adaptors are the same length (Figure S7B).

### Functions of RH1 domain-containing proteins other than JIP3

JIP3 is part of a family of RH1 domain-containing proteins^92,93^. Its closest homolog is JIP4, which is associated with lysosome movement in non-neuronal cells^94^. The dynein-dynactin interacting regions are highly conserved between JIP3 and JIP4 suggesting both work in a similar way. A more distant relative is RILP, which binds the small G-protein Rab7^95^, recruits dynein-dynactin to late endosomes and lysosomes^96^ and interacts with the DLIC C-terminus^37,97^. RILP’s N-terminal coiled coil is a similar length to JIP3’s LZI, although how it binds dynein-dynactin is not clear. Unlike for JIP3 and JIP4, AlphaFold2 cannot predict an HBS1 interaction with the DHC. Furthermore, although RILP contains a potential Spindly motif (LLLEA - residues 284-289), this is C-terminal to the cargo binding (Rab7) site, compared to all known adaptors where the Spindly motif is on the N-terminal side.

Other family members include RILPL1 and RILPL2. Of these, RILPL1 can bind the DLIC C-terminus^37^ and can cluster objects at the center of the cell^92^ consistent with a role in dynein-dynactin function. However, AlphaFold2 again does not predict an HBS1 interaction and there is no obvious Spindly motif. RILPL2 does not bind dynein and is instead implicated in Myosin-Va binding^98^. Interestingly, all the RH1 domain family members, even RILPL2, contain helical segments analogous to the JIP3^helix^ (Figure S7C). This suggests the autoinhibition mechanism observed in JIP3 is likely to be conserved across this protein family.

### Role of p150 and DIC-N in DDA assembly

Our cryo-EM structure shows the dynactin p150 arm in its open conformation making contacts with dynein and LIS1. We also used AlphaFold2 and previously published cryo-EM data to produce a molecular model of p150’s inhibited state. A comparison of these structures shows that dynactin autoinhibition involves both the previously identified hairpin formed by the CC1A binding to CC1B^38,40,49^ and an additional interaction of p150’s globular ICD domain. Both CC1A and ICD need to detach for CC1B to bind dynein’s motor and LIS1. In addition, opening up p150 allows the ICD to contact dynein’s tail. Our observations implicate the ICD, which previously had no known function, as part of the mechanism of dynactin autoinhibition and DDA assembly.

The open p150 binds to two copies of the DIC-N region from dynein-A. Surprisingly, in the inhibited form most of the two DIC-N binding sites on CC1B are accessible. Only the edges of the sites are partially occluded by the ICD and CC1A. This explains how DIC-N opens up p150 by displacing both these regions. The fact that the DIC-N binding region remains partly accessible explains many pull-down studies showing an interaction between DIC-N and p150^41–46,48^. Curiously, other studies using gel-filtration^13,14^, sucrose gradients^56^ or single molecule assays^12^ reported limited interaction of dynein and dynactin unless an activating adaptor is present. This may reflect the tendency of these methods to detect higher affinity interactions than pull-downs, suggesting that although dynein and dynactin can interact though DIC-N:p150, the formation of DDA adds further stability to this interaction.

It was previously not entirely clear why DDA complex formation requires the binding of DIC-N to CC1B^48^. Our structure shows that in comparison to DIC-N, the binding of LIS1 and dynein’s motor domain to CC1B is incompatible with the inhibited form of p150 (Figure 4). In addition, biochemical studies show that whereas DIC-N can bind p150 on its own, the interaction of LIS1 requires the presence DIC-N (Figure S4). Together these results suggest that a key reason DIC-N binds CC1B is to open up p150 to allow subsequent interactions with LIS1 and the dynein motor to occur.

### The IC-LC tower facilitates the p150 – DIC interaction during transport

In our structure, the IC-LC towers of both dynein-A and dynein-B are docked under their respective heavy chains (Figure S5). As the two dyneins differ in both their conformation and the presence of LIS1, it suggests neither attribute is required for IC-LC docking. We also looked back at our previous dynein-dynactin-BICDR1 (DDR) structure on microtubules^27^, which was prepared without LIS1, and found evidence for light chains in a similar position. Together these observations suggest that the standard position for dynein light chains is underneath the DDA complex. This would make it unlikely they play a direct role in binding cargos, in contrast to a number of previous studies^99–101^, but in agreement with others^81,102^.

Studies on isolated dynein show the IC-LC tower either extending away from the motor^39^ or docked onto one heavy chain via the LC8 light chain^14^. In our structure this LC8 interaction is the same^14^ whereas the additional TCTEX1 interaction is only possible because of the parallel arrangement of the heavy chains. This additional interaction could help stabilise the parallel arrangement of dynein motors needed for processive movement^14^. Interestingly, light chain – heavy chain interactions are also found in dynein-2^103^ and axonemal dyneins^104–108^. However, in all these cases the light chains make only a single contact with the relevant heavy chain, even in those examples where the heavy chains are parallel^104–107^. This raises the possibility that the additional light chain – heavy chain interactions observed in our study are specific to DDA complexes.

The position of the IC-LC tower on dynein-A constrains the DIC-Ns, and in turn p150, to be close to the dynein-A motor domains. Furthermore, a recent single molecule study suggested that DIC-N remains bound to p150 during dynein-dynactin movement^48^. These observations explain the position of an extra density, similar in shape to p150, that we reported in our previous DDR structure^27^. There are a number of implications for positioning p150 close to dynein within a moving DDA complex. Firstly, it orients p150 such that its microtubule binding segments are held close to the microtubule (Figure S7D), consistent with its role in increasing dynein’s interaction with microtubules^74,75,109–111^. Secondly, it means the p150 is available to interact with dynein-A1 whenever it is in its pre-powerstroke conformation. Continual binding/unbinding would be expected to interfere with the ATP hydrolysis cycle of dynein. This effect would be more consequential in single-dynein DDA complexes and may explain their slower stepping rate and velocity compared to two-dynein DDA complexes^70,112^.

### LIS1 keeps dynein in a low microtubule affinity state

A striking feature of our structure is that the two dynein motors which bind LIS1 (dynein-A1/A2) are in a pre-powerstroke conformation with their stalks raised up and detached from the microtubule. Several previous studies showed LIS1 preferentially binds dynein motors in their pre-powerstroke conformation^64–67,79^. Here, our structure was solved in the presence of AMP-PNP which strongly favours post-powerstroke dynein. This suggests LIS1 doesn’t just recognise the pre-powerstroke state of dynein, but rather induces it by overriding the conformation dictated by the nucleotide. This agrees with a previous study^67^, although in that case it was reported that ATP/ADP-Pi needed to be present in AAA3 for LIS1 to induce a pre-powerstroke state. In contrast, our structure suggests ATP/ADP-Pi in AAA3 is not absolutely required since we see density for ADP in AAA3. This difference might be due to additional interactions within our DDA complex that were not present in truncated motor domain constructs used previously. These include LIS1 simultaneously contacting the two dynein-A motors and the attached p150.

In our structure, dynein-B is in a post-powerstroke conformation and lacks LIS1. Given reports that LIS1 encourages binding of a second dynein dimer^59,60^ it seems probable that dynein-B also engages LIS1 during DDA complex assembly. Unlike the dynein-A bound LIS1 molecules, which are additionally stabilised by the presence of p150, the LIS1 molecules on dynein-B are likely to have been displaced upon productive microtubule binding^79,83^. Dynein-B will thus be the first motor to contact the microtubule. There is conflicting evidence on the extent to which LIS1 can associate with motile DDA complexes, however all groups report some colocalization during transport^59–61,69^. Our structure provides an explanation for how LIS1 can remain attached to moving DDA complexes via dynein-A and p150, while the movement is driven by dynein-B.

### Interaction of LIS1 with p150 is important for DDA assembly

Although many roles for LIS1 have been proposed^17,113^, recent data suggested its major function is stimulating DDA complex assembly^59–61,69^. The current model suggests LIS1 achieves this by binding dynein motor domains and opening up the inhibited phi-particle^17,62,63,68^. However, we find that LIS1 constructs which bind dynein, but not p150, are unable to stimulate DDA complex formation *in vitro* or rescue loss of LIS1 in cells (Figure 6). This suggests that opening the phi-particle, while likely important, is not sufficient to promote DDA assembly. Instead, the ability of LIS1 to contact both p150 and dynein is required.

Why might the LIS1 – p150 interaction be needed for the formation of active DDA complex? Firstly, our structure suggests LIS1 reinforces the tethering of dynein to p150. Once bound to CC1B, LIS1 would help keep the CC1A/B hairpin in its open conformation favouring strong DIC-N binding. This would also explain LIS1’s ability to enhance the recruitment of dynein to dynactin at microtubule plus ends^56,61,69^.

Secondly, our structure provides evidence that LIS1 orients dynein-A for optimal dynactin and adaptor binding. The ability of LIS1 to contact both the CC1B and dynein allows p150 to dock along the heavy chain by binding the dynein tail and motor. We propose this tucks dynein under the p150 arm and holds its tail next to the dynactin filament (Figure 7). This would restrict the freedom of dynein and dynactin to prime them for efficient conversion into a motile complex in the presence of an activating adaptor.

**Figure 7.**
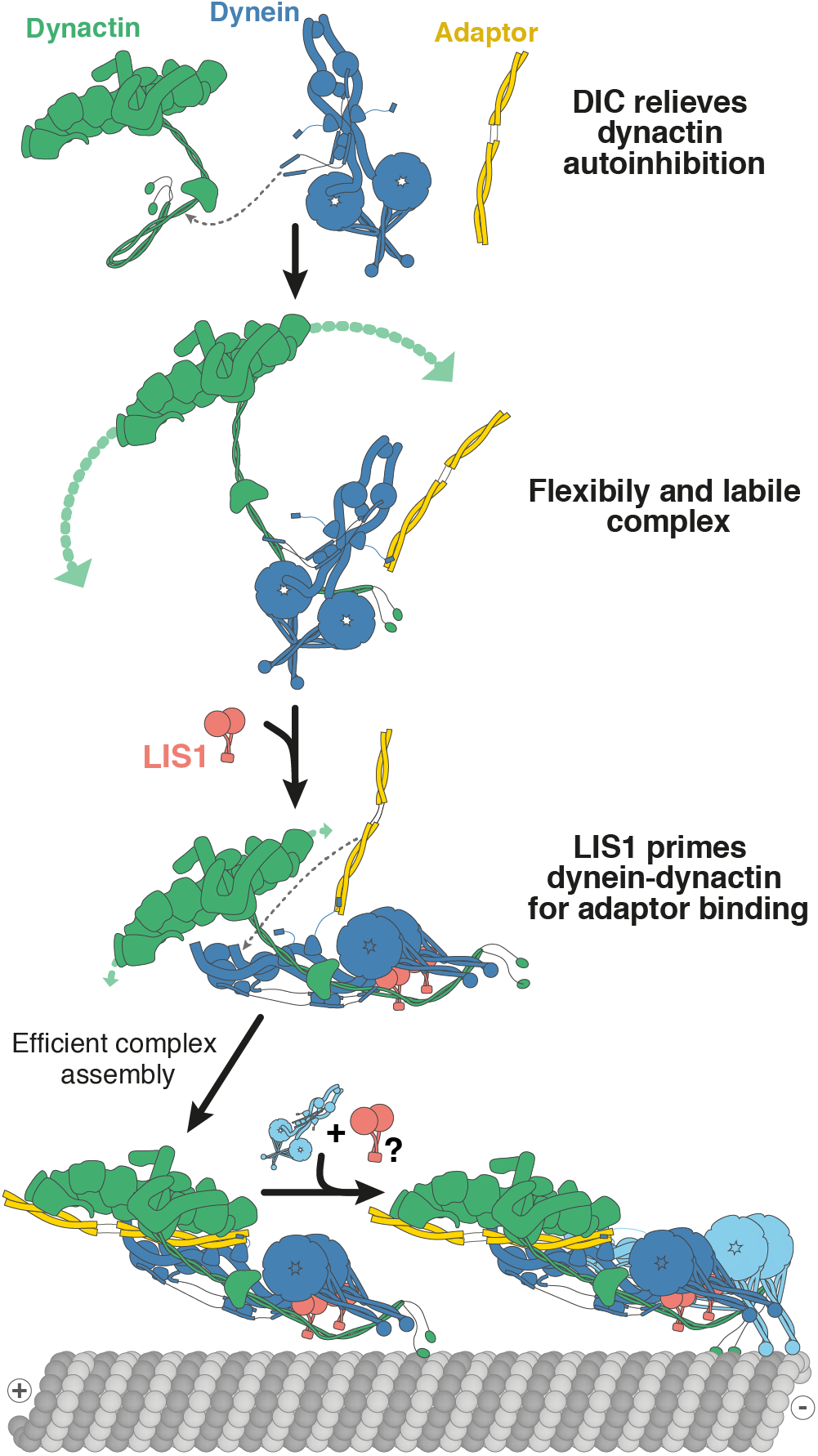
Model for formation of DDA complexes. We propose a model for how DDA complexes assemble in the presence of LIS1. The binding DIC to p150 helps relieve dynactin autoinhibition by promoting the open form of the p150 arm. However, this interaction may be labile and flexible. The binding of LIS1 to p150 will then reinforce the interaction between DIC-N and p150. In addition, LIS1 stabilises the pre-powerstroke state of dynein allowing for the p150 to dock along dynein as well as directly tethers p150 to a dynein motor. The binding of LIS1 to the motor domains would also promote a parallel dynein arrangement which is required for the dynein tails to bind to the dynactin filament. Through their interactions, LIS1 and p150 work together to orient the dynein tails near the dynactin filament and prime the complex for efficient conversion into a motile complex in the presence of an activating adaptor in its open form. The next step is the recruitment of another dynein molecule which will be the first to bind microtubules in two dynein complexes.

### Model for formation of DDA complexes

The data presented in this manuscript provides evidence for a number of steps in the formation of DDA complexes (Figure 7). We show that dynactin’s p150 arm is autoinhibited by both a CC1A/B hairpin and interactions with the ICD domain. Dynein binding via DIC-N appears to be required first and its role is to open up the p150. This in turn allows formation of a complex containing interactions between p150, LIS1, the dynein motor and the dynein tail. We assume at this stage the cargo-bound activating adaptor is already open and bound to dynein via the DLIC interaction, although, it is also possible dynein-dynactin-LIS1 complexes may form without the adaptor present. In either case, the adaptor must next move into its final position, allowing the dynein tails to fully interact with dynactin. For complexes that contain two dyneins, the next step is recruitment of dynein-B, which our structure suggests can readily interact with the microtubule, leading to the initiation of processive movement.

## Methods

### Constructs

Full-length human JIP3 sequence (Uniprot Q9UPT6-1) was codon optimised for expression in Sf9 cells (Twist Bioscience). Residues 1-1336 (JIP3FL), 1-582 (JIP31-582), 1-560 (JIP31-560) were subcloned with a N-terminal His6-ZZ tag followed by a TEV protease cleavage site (TEV) in a pAceBac1 vector. JIP31-582_F576A/F577A and JIP31- 560_L382A/Y383A/E385A were generated using site directed mutagenesis of JIP31-582 and JIP31-560 constructs. Residues 1-560 were also inserted into a 2CT vector [N-terminal 6xHis::maltose binding protein (MBP) followed by a TEV protease cleavage site and C-terminal Strep-tag II (StTgII)] for bacterial expression. Constructs coding for JIP3 residues 1-108, 1-185 and 186-50537 were constructed by cloning fragments from mouse cDNA (isoform 3a; Uniprot Q9ESN9-5) into the 2CT vector mentioned above. For the N-terminal JIP3 fragments, the only difference between the mouse and human sequence is residue 102 (lysine versus arginine, respectively). Mouse JIP3 186-505 differs from the equivalent human fragment at multiple residues, but the Spindly motif and the Arf6 binding site are identical. Fragments coding for human JIP3helix (563-585) and human Arf6 (12-175_Q67L) were inserted into a pGEX-6P-1 vector [N-terminal glutathione S-transferase (GST) followed by a Prescission protease (Psc) cleavage site and C-terminal 6xHis]. JIP3 mutants 1- 108_V60Q37, 186-505_L439P37, 186-505_L383A/E386A, and 563-585_F576A/L579A were generated using site directed mutagenesis.

For the HOOK3 adaptor, pACEBac1-2xStrep-Psc-HOOK31–522 was used and purified as previously described^70^.

The dynactin pointed end complex construct28 comprised of human ZZ-TEV-Arp11, p62 (isoform A, UniProt Q9UJW0-1), p25 and p27 in a pAceBac1 vector was used in the gel-filtration experiment (Figure S3C). It was purified as described previously28. For pulldown experiments, the pointed end complex was prepared as described29 and was comprised of Arp11, p62, p27 and p25 tagged with a C-terminal 6xHis tag.

For dynein, full-length wild-type human cytoplasmic dynein-113 and a “phi” mutant construct with mutations in the linker (R1567E and K1610E) to help overcome the autoinhibited conformation14 were used. Dynein intermediate chain sequence 1-76 was cloned into the 2CT vector mentioned above. GST-Psc-DCTN1(residues 214-547)-GFP-6xHis, GST-Psc-DCTN1(residues 214-354)-GFP-6xHis and GST-Psc-DCTN1(residues 350-547)-GFP-6xHis were used for CC1, CC1A and CC1B fragment of DCTN1/p150glued (Uniprot Q14203-1), respectively.

Full length human LIS1 with an N-terminal ZZ-TEV tag or a N-terminal ZZ-TEV tag plus a C-terminal SNAPf tag was cloned into a pFastbac vector61. These constructs were used to generate LIS1 truncations - LIS1-N (residues 1-90), WD40 (residues 90-410) and GCN4-WD40 (GSGSVKQLEDKVEELLSKNAHLENEVARLKKLV + LIS1_81-410). For constructing LIS1mut (LIS1_L3A_R6E_Q7A_K46E_K53E_K54E_T56A_S57A_I5 9A_R60E_Q62A_K63E_K64E_E69R), LIS1-N-FKBP (LIS1_1-84::5X-GS-linker::FKBP) and FRB-WD40 (FRB-T2098L::5X-GS-linker::LIS1_91-410), the sequences were first synthesised (Twist Bioscience) and then inserted into the pFastbac vector.

### Protein expression

For bacterial expression of proteins, SoluBL21 *E. coli* cells (Genlantis) were used and the expression was induced overnight at 18°C with 0.1-0.25 mM IPTG added at O.D of 0.6-0.8. Cells were harvested by centrifugation at 4,000 rcf for 20 min followed by a PBS wash and another centrifugation at 4,000 rcf for 20 min. The cell pellet was flash frozen in liquid nitrogen and stored at -80°C.

For expression in Sf9 cells, fresh bacmid DNA was transfected into Sf9 cells at 0.5×10^6^ cells/mL in 6-well cell culture plates using FuGene HD (Promega) according to the manufacturer’s protocol (final concentration 10 μg/mL). After six days, 1 mL of the culture supernatant was added to 50 mL of 1×10^6^ cells/mL and cells were infected for five days in a shaking incubator at 27°C. The P2 virus was isolated by collecting the supernatant after centrifugation at 4,000 rcf for 15 min and stored at 4°C. For expression, 10 mL of P2 virus was used to infect 1 L of Sf9 cells at 1.5-2×10^6^ cells/mL for 72 hours in a shaking incubator at 27°C. Cells were harvested by centrifugation at 4,000 rcf for 10 min at 4°C, and washed with cold PBS. The cell pellet was flash frozen and stored at -80°C.

### Protein purification

Dynactin was purified from frozen porcine brains as previously described^40^. Fresh brains were cleaned in homogenization buffer (35 mM PIPES pH 7.2, 5 mM MgSO4, 100 μM EGTA, 50 μM EDTA), and flash frozen in liquid nitrogen. Frozen brains were broken into pieces using a hammer. The brain pieces were blended and resuspended in homogenization buffer supplemented with 1.6 mM PMSF, 1 mM DTT, and 4 complete-EDTA protease-inhibitor tablets (Roche) per 500 mL. After thawing, the lysate was centrifuged in a JLA 16.250 rotor (Beckman Coulter) at 16,000 rpm for 15 min at 4°C. The supernatant was further clarified in a Type 45 Ti rotor (Beckman Coulter) at 45,000 rpm for 50 min at 4°C. After filtering the supernatant in a Glass Fibre filter (Sartorius) and a 0.45 μm filter (Elkay Labs), it was loaded on a column packed with 250 mL of SP-Sepharose (Cytiva) pre-equilibrated with SP buffer (35 mM PIPES pH 7.2, 5 mM MgSO4, 1 mM EGTA, 0.5 mM EDTA, 1 mM DTT, 0.1 mM ATP) using an Akta Pure system (Cytiva). The column was washed with SP buffer with 3 mM KCl before being eluted in a linear gradient up to 250 mM KCl over 3 column volumes. The peak around ∼15 mS/cm was collected and filtered in a 0.22 μm filter (Elkay Labs) before being loaded on a MonoQ 16/10 column (Cytiva) pre-equilibrated with MonoQ buffer (35 mM PIPES pH 7.2, 5 mM MgSO4, 100 μM EGTA, 50 μM EDTA, 1 mM DTT). The column was washed with MonoQ buffer before being eluted in a linear gradient up to 150 mM KCl over 1 column volume, followed by another linear gradient up to 350 mM KCl over 10 column volumes. The peak around ∼39 mS/cm was pooled and concentrated to ∼3 mg/mL before being loaded on a TSKgel G4000SWXL column (Tosoh Bioscience) preequilibrated with GF150 buffer (25 mM HEPES pH 7.2, 150 mM KCl, 1 mM MgCl2) supplemented with 5 mM DTT and 0.1 mM ATP. The peak at ∼114 mL was pooled and concentrated to ∼3 mg/mL. 3 μL aliquots were flash frozen in liquid nitrogen and stored at -80°C.

For dynein purification, a cell pellet from 1 L expression was resuspended in 50 mL lysis buffer (50 mM HEPES pH 7.4, 100 mM NaCl, 10% (v/v) glycerol, 0.1 mM ATP) supplemented with 2 mM PMSF, 1 mM DTT, and 1 complete-EDTA protease-inhibitor tablet. Cells were lysed using a 40 mL dounce tissue grinder (Wheaton) with ∼20 strokes. The lysate was clarified at 503,000 rcf for 45 min at 4°C using a Type 70 Ti Rotor (Beckman Coulter). The supernatant was incubated with 3 mL IgG Sepharose 6 Fast Flow beads (Cytiva) pre-equilibrated with lysis buffer for 4 hours at 4°C. The beads were then applied to a gravity flow column and washed with 150 mL of lysis buffer and 150 mL of TEV buffer (50 mM Tris-HCl pH 7.4, 150 mM KAc, 2 mM MgAc, 1 mM EGTA, 10% (v/v) glycerol, 0.1 mM ATP, 1 mM DTT). For TMR labelled dynein, beads were transferred to a tube and incubated with 10 μM SNAP-Cell TMR-Star dye (New England Biolabs) for 1 hour at 4°C prior to the TEV buffer washing step. The beads were then transferred to a 5 mL centrifuge tube (Eppendorf) and filled up completely with TEV buffer. 400 μg TEV protease was added to the beads followed by overnight incubation at 4°C. The beads were transferred to a gravity flow column and the flow through containing the cleaved protein was collected. The protein was concentrated to ∼2 mg/mL and loaded onto a TSKgel G4000SWXL column pre-equilibrated with GF150 buffer supplemented with 5 mM DTT and 0.1 mM ATP. Peak fractions were pooled and concentrated to ∼2.5-3 mg/mL. Glycerol was added to a final concentration of 10% from an 80% stock made in GF150 buffer. 3 μL aliquots were flash frozen and stored at -80°C.

For purification of LIS1 constructs (full-length, truncations and mutants), a cell pellet from 1 L expression was resuspended in 50 mL lysis buffer B (50 mM Tris-HCl pH 8.0, 250 mM KAc, 2 mM MgAc, 1 mM EGTA, 10% (v/v) glycerol, 0.1 mM ATP, 1 mM DTT) supplemented with 2 mM PMSF. Cells were lysed using a 40 mL dounce tissue grinder (Wheaton) with ∼20 strokes. The lysate was clarified at 150,000 rcf for 30 min at 4°C using a Type 45 Ti Rotor (Beckman Coulter). The supernatant was incubated with 3 mL IgG Sepharose 6 Fast Flow beads (Cytiva) pre-equilibrated with lysis buffer B for 4 hours at 4°C. The beads were then applied to a gravity flow column and washed with 150 mL of lysis buffer B. The beads were then transferred to a 5 mL centrifuge tube (Eppendorf) and filled up completely with lysis buffer B. 400 μg TEV protease was added to the beads followed by overnight incubation at 4°C. The beads were transferred to a gravity flow column and the flow through containing the cleaved protein was collected. The protein was concentrated to ∼3-5 mg/mL and loaded onto a Superdex 200 Increase 10/300 (Cytiva) column pre-equilibrated with GF150 buffer supplemented with 5 mM DTT and 10% glycerol. Peak fractions were pooled and concentrated to ∼2-5 mg/mL. 5 μL aliquots were flash frozen and stored at -80°C.

For JIP3 constructs expressed in Sf9 cells, cell pellet from 1 L expression was resuspended in 50 mL lysis buffer C (50 mM HEPES pH 7.5, 100 mM NaCl, 10% (v/v) glycerol) supplemented with 2 mM PMSF, 1 mM DTT, and 1 complete-EDTA protease-inhibitor tablet. Cells were lysed using a 40 mL dounce tissue grinder (Wheaton) with ∼20 strokes. The lysate was clarified at 150,000 rcf for 45 min at 4°C using a Type 45 Ti Rotor (Beckman Coulter). The supernatant was incubated with 3 mL IgG Sepharose 6 Fast Flow beads (Cytiva) pre-equilibrated with lysis buffer for 4 hours at 4°C. The beads were then applied to a gravity flow column and washed with 150 mL of lysis buffer C and 150 mL of TEV buffer B (50 mM Tris-HCl pH 7.4, 150 mM KAc, 2 mM MgAc, 1 mM EGTA, 10% (v/v) glycerol, 0.1 mM ATP). The beads were then transferred to a 5 mL centrifuge tube (Eppendorf) and filled up completely with TEV buffer B. 400 μg TEV protease was added to the beads followed by overnight incubation at 4°C. The beads were transferred to a gravity flow column and the flow through containing the cleaved protein was collected. The protein was concentrated to ∼2-4 mg/mL and loaded on a Superose 6 Increase 10/300 column (Cytiva) equilibrated with storage buffer (25 mM HEPES pH 7.5, 150 mM NaCl, 1 mM MgCl2, 1 mM DTT). Glycerol was added to a final concentration of 10% (v/v) and aliquots were flash frozen in liquid nitrogen and stored at - 80°C.

For bacterially expressed MBP-StTgII tagged constructs, cell pellet from a 2 L culture was resuspended in 50 mL lysis buffer D (50 mM HEPES pH 8.0, 250 mM NaCl, 10 mM imidazole, 0.1% Tween 20, 1 mM DTT, 1 mM PMSF, 2 mM benzamidine-HCl) supplemented with 1 complete-EDTA protease-inhibitor tablet and 1 mg/mL Lysozyme. Cells were lysed by sonication and the lysate was clarified at 150,000 rcf for 30 min at 4°C using a Type 45 Ti Rotor (Beckman Coulter). The lysate was passed three times though 2mL of pre-equilibrated Ni-NTA Agarose beads (Qiagen) in a gravity flow column and then washed with 300mL of wash buffer A (25 mM HEPES pH 8.0, 250 mM NaCl, 25 mM imidazole, 0.1% Tween 20, 1 mM DTT, 1 mM PMSF, 2 mM benzamidine-HCl). Proteins were eluted with elution buffer A (50 mM HEPES pH 8.0, 150 mM NaCl, 250 mM imidazole, 1 mM DTT, 2 mM benzamidine-Cl). Fractions containing the protein were pooled, incubated overnight at 4°C with TEV protease and then incubated in batch with Strep-Tactin Sepharose resin (IBA) for 1 hour at 4°C, and washed with wash buffer B (25 mM HEPES pH 8.0, 250 mM NaCl, 0.1 % Tween 20, 1 mM DTT). Proteins were eluted on a gravity column with elution buffer B (50 mM HEPES pH 7.5, 150 mM NaCl, 3.5 mM desthiobiotin). The protein was further purified by size exclusion chromatography on a Superose 6 Increase 10/300 column (Cytiva) or Superdex 200 Increase 10/300 GL column (Cytiva) equilibrated with storage buffer (25 mM HEPES pH 7.5, 150 mM NaCl, 1 mM MgCl2, 1 mM DTT). Glycerol was added to a final concentration of 10% (v/v) and aliquots were flash frozen in liquid nitrogen and stored at - 80°C.

For bacterially expressed GST-6xHis tagged constructs, cell pellet from a 2 L culture was resuspended in 50mL lysis buffer E (20 mM Tris-HCL pH 8.0, 300 mM NaCl, 15 mM imidazole, 10% glycerol, 1 mM DTT, 1 mM PMSF) supplemented with 1 complete-EDTA protease-inhibitor tablet and 1 mg/mL Lysozyme. Cells were lysed by sonication and the lysate was clarified at 150,000 rcf for 30 min at 4°C using a Type 45 Ti Rotor (Beckman Coulter). The lysate was passed three times through 2 mL of pre-equilibrated Ni-NTA Agarose beads (Qiagen) in a gravity flow column and then washed with 300 mL of lysis buffer D. Proteins were eluted elution buffer C (20 mM Tris-HCL pH 8.0, 300 mM NaCl, 200 mM imidazole, 10% glycerol, 1 mM DTT). Fractions containing the protein were pooled and then incubated in batch with glutathione agarose resin (Thermo Fisher Scientific) for 1 hour at 4°C and washed with wash buffer B (20 mM Tris-HCL pH 8.0, 300 mM NaCl, 1 mM DTT). Proteins were eluted on a gravity column with elution buffer D (20 mM Tris-HCL pH 8.0, 300 mM NaCl, 20 mM reduced glutathione, 1 mM DTT). The protein was further purified by size exclusion chromatography on a Superose 6 Increase 10/300 column (Cytiva) equilibrated with GF150 (25 mM HEPES pH 7.5, 150 mM KCl, 1 mM MgCl2, 1 mM DTT). Glycerol was added to a final concentration of 10% (v/v) and aliquots were flash frozen in liquid nitrogen and stored at - 80°C.

### Pull-down assays

For strep-tagged proteins, purified recombinant proteins (50 - 300 pmol each) were mixed in a total of 20 μL Strep-PD buffer (25 mM HEPES pH 7.5, 150 mM NaCl, 1 mM DTT) and incubated at room temperature for 30 min. For GST-and MBP-tagged proteins, GST-PD buffer (50 mM HEPES pH 7.5, 100 mM KCl, 1mM MgCl2, 1 mM DTT) was used. 4 μL were removed from the mixture ("input") before adding 30 μL of a 50 % resin slurry (Glutathione Sepharose (Cytiva) for GST pull-downs, Strep-Tactin Sepharose (IBA Lifesciences) for Strep-tag II pull-downs and Amylose resin (NEB) for MBP pull-downs). The resin/protein mixture was rotated at room temperature for 30 min, and the resin was washed with 3 x 500 μL of the respective PD buffer, which for Strep-tag II pull-downs was supplemented with 0.05 % Tween-20. Proteins were eluted with 40 μL of 10 mM reduced L-glutathione in GST-PD buffer (glutathione agarose resin), 10 mM Maltose in GST-PD buffer (Amylose resin) or with 10 mM *d*- desthiobiotin in 100 mM Tris-HCL pH 8, 150 mM NaCl, 1 mM EDTA (Strep-Tactin Sepharose resin). 23 μL of PD buffer and 9 μL of 4x SDS-PAGE sample buffer were added to 4 μL of the input, and 9 μL of 4x SDS-PAGE sample buffer were added to 27 μL of the eluate before incubating samples at 95°C for 1 min. 5 μL of the input sample and 10 μL of the eluted sample were separated by SDS-PAGE and visualized by Coomassie Blue staining.

### Gel filtration for JIP3-dynactin pointed end binding

Dynein pointed end complex and JIP3^1–560^ or JIP3^1–560^ with mutations in the Spindly motif (L382A_Y383A_E385A) were mixed in 1:1 ratio in GF150 buffer with a final concentration of 10 μM in a volume of 60 μL. The mix was incubated for 30 min at 4°C and then 50 μL were loaded on a Superose 6 Increase 3.2/300 column (Cytiva) equilibrated with GF150.

### *In vitro* TIRF motility assays

Microtubules were made by mixing 1.5 μL of 2 mg/mL HiLyte Fluor 488 tubulin (Cytoskeleton), 2 μL of 2 mg/mL, biotinylated tubulin (Cytoskeleton) and 6.5 μL of 13 mg/mL unlabelled pig tubulin^13^ in BRB80 buffer (80 mM PIPES pH 6.8, 1 mM MgCl2, 1 mM EGTA, 1 mM DTT). 10 μL of polymerization buffer (2× BRB80 buffer, 20% (v/v) DMSO, 2 mM Mg-GTP) was added followed by incubation at 4°C for 5 min. Microtubules were polymerized at 37°C for 1 h. The sample was diluted with 100 μL of MT buffer (BRB80 supplemented with 40 μM paclitaxel), then centrifuged on a benchtop centrifuge (Eppendorf) at 21,000 rcf for 9 minutes at room temperature. The resulting pellet was gently resuspended in 100 μL of MT buffer, then centrifuged again as above. 50 μL MT buffer was then added and the microtubule solution was protected from light. Microtubules were prepared the day before the assay was performed and stored in pellet form. Before usage, and every 5 hours during data collection, the microtubule solution was spun again at 21,000 rcf for 9 minutes, and the pellet resuspended in the equivalent amount of MT buffer.

Motility chambers were prepared by applying two strips of double-sided tape approximately 8-10 mm apart on a glass slide and then placing a piranha-solution-cleaned coverslip on top. The coverslip was functionalized using PLL-PEG-Biotin (SuSOS AG), washed with 50 μL of TIRF buffer (30 mM HEPES pH 7.2, 5 mM MgSO4, 1 mM EGTA, 2 mM DTT), then incubated with streptavidin (1 mg/mL, New England Biolabs). The chamber was again washed with TIRF buffer, then incubated with 10 μL of a fresh dilution of microtubules (1.5 μL of microtubules diluted into 10 μL TIRF-Casein buffer (TIRF buffer supplemented with 50 mM KCl and 1 mg/mL casein) for 1 min. Chambers were then blocked with 50 μL TIRF-Casein buffer.

Complexes were prepared by mixing each component in a total volume of 6 uL in GF150. The final concentrations were TMR-dynein at 0.1 μM, dynactin at 0.2 μM, adaptor at 2 μM and LIS1 at 6 μM. For LIS1 constructs where two components were added (for example: GCN4-WD40 + LIS1- N) each component was added at a final concentration of 6 μM in the initial mix. Complexes were incubated on ice for 15 min then diluted with TIRF-Casein buffer containing 75 mM KCL to a final volume of 10 μL. 1 μL of this complex was added to 19 μL of TIRF-Casein buffer supplemented with an oxygen scavenging system (0.2 mg/mL catalase, Merck; 1.5 mg/mL glucose oxidase, Merck; 0.45% (w/v) glucose), 1% BME, 5 mM Mg-ATP. This mix was flowed into the chamber. The sample was imaged immediately at 23°C using a TIRF microscope (Nikon Eclipse Ti inverted microscope equipped with a Nikon 100x TIRF oil immersion objective). For each sample, a microtubule image was acquired using a 488 nm laser. Following this a 500-frame movie was acquired (200 ms exposure, 4.1 fps) using a 561 nm laser. To analyse the data, ImageJ was used to generate kymographs from the tiff movie stacks. Events of similar length were picked to analyse velocity, run length and number of processive events/μm microtubule/min, using criteria outlined previously^13,70^. Velocity was calculated using pixel size of 105 nm and frame rate of 236 ms/frame. Three replicates were performed for each sample. Statistical significance was determined using ANOVA with Tukey’s multiple comparison test.

### Cryo-EM sample preparation

The sample was prepared in a similar manner as described previously^27^. To polymerize microtubules, tubulin was diluted in microtubule buffer (25 mM MES pH 6.5, 70 mM NaCl, 1 mM MgCl2, 1 mM EGTA, 1 mM DTT) with 6 mM GTP (MilliporeSigma) such that the final concentration of tubulin was 5 mg/mL (45 μM) tubulin and that of GTP was 3 mM. The mixture was incubated on ice for 5 min, and microtubules were polymerized at 37°C for ∼1.5 hours. To stabilize the microtubules, polymerization buffer was supplemented with 20 μM paclitaxel. The microtubules were pelleted on a benchtop centrifuge (Eppendorf) at 20,000 rcf for 8 min at room temperature. The supernatant was discarded and the pellet was resuspended in polymerization buffer with paclitaxel by pipetting up and down with a cut tip. The microtubules were pelleted and resuspended again using an uncut tip. The concentration was estimated using Bradford reagent (BioRad) and diluted to ∼0.65 mg/mL (∼6 μM). To assemble the dynein-dynactin-JIP3 complex in the presence of LIS1, the purified proteins were mixed in a 1:2:60/44:32 molar ratio (0.27 μM dynein, 0.54 μM dynactin, 16 μM JIP3^1–185^ or 12 μM for JIP3^1–560^ and 8.7 μM LIS1) in GF150 buffer supplemented with 1 mM DTT in a total volume of 10 μL and incubated on ice for 30 min. To bind the complex to microtubules, 9 μL complex was mixed at room temperature with 5 μL microtubules and 9 μL binding buffer A (77 mM HEPES pH 7.2, 51 mM KCl, 13 mM MgSO4, 2.6 mM EGTA, 2.6 mM DTT, 7.68 mM AMPPNP, 13 μM paclitaxel) such that the final concentrations of KCl and AMPPNP were 100 mM and 3 mM, respectively. After 15 min, the complex-bound microtubules were pelleted at 20,000 rcf for 8 min at room temperature. The pellet was resuspended in binding buffer B (30 mM HEPES pH 7.2, 50 mM KCl, 5 mM MgSO4, 1 mM EGTA, 1 mM DTT, 3 mM AMPPNP, 5 μM paclitaxel, and 0.01% IGEPAL (MilliporeSigma) using an uncut tip. 3.5 μL was applied to freshly glow discharged Quantifoil R2/2 300-square-mesh gold grids (Quantifoil) in a Vitrobot IV (ThermoFisher) at 100% humidity and 20°C, incubated for 30 s, and blotted for 0.5-2 s before being plunged into liquid ethane.

### Cryo-EM data collection and image processing

The samples were imaged using a FEI Titan Krios (300 kV) equipped with a K3 detector and energy filter (20 eV slit size) (Gatan) using automated data collection (ThermoFisher EPU). Movies were acquired at 81,000 X magnification (1.06 Å/pixel, 100 μm objective aperture, ∼2.2 sec exposure, 54 frames, fluence of ∼54 e-/Å^2^ and -1.2 to -3 μm defocus range).

Global motion correction and dose-weighting were performed in Relion-4.0^115^ using MotionCorr2^116^ with a B-factor of 150 and 5X5 patches. The power spectrum of aligned frames was used for CTF estimation by CTFFIND4^117^. Microtubules were picked from the micrographs using the single particle picking mode in crYOLO^118^ with a model that was trained on manually picked micrographs (∼100) in filament mode. The output coordinates along each microtubule were spaced by 81 Å. The coordinates were then resampled to a 4 nm interval using the multi-curve fitting script described previously^72^ (https://github.com/PengxinChai). The coordinates for each microtubule were then further split into segments of 10-17 coordinates as this gave better results with the following microtubule subtraction step. Finally, microtubule subtraction was performed as described previously^72^. The subtracted microtubules were then used to pick dynein-dynactin-JIP3 complexes using crYOLO^119^. The picked particles were imported into CryoSPARC^120^ followed by 2D classification. The particles in classes showing dynein-dynactin were used for ab-initio initial model generation. Three initial models were generated which were then used for a round of heterogeneous refinement using all particles. Particles belonging to the 3D class which displayed good density for dynein-dynactin were used for a round of non-uniform refinement and then imported into Relion-4.0 for particle polishing and local refinements. The consensus structures of complexes with JIP3^1–185^ and JIP3^1–560^ appeared identical except the JIP3 densities at the pointed end in the DDLJ^1–560^ dataset. Therefore, the two datasets were merged at this stage. The merged data was used for local refinements and classifications for all parts of the complex except the dynactin pointed end which was solved using only the DDLJ^1–560^ dataset. The processing after this stage was performed in a similar manner as described before^27^. The processing pipeline including mask used, particle numbers, resolution of maps, B-factors and classification parameters can be found in Figure S2. Data collection and image processing statistics can be found in Table S2.

### Model building and refinement

Model building and restrained flexible fitting of models in the density was done in COOT^121^ and refinements were performed in PHENIX^122^. Models from the Protein Data Bank (PDB) or AlphaFold2 predictions (see section below) were docked into the respective cryo-EM maps as a rigid body using UCSF Chimera^123^. All refinement statistics can be found in Table S2.

AlphaFold2 predictions of JIP3 (1-185) and the JIP3 RH1 domain bound to DLIC^helix^ were used as starting models for JIP3 and DLIC^helix^. The registry of the JIP3 coiled-coil was determined using the characteristic shape of the RH1 domain and side chain resolution at the HBS1 site.

Dynein tails with bound dynein subunits (DLIC RAS-like domain, C-terminal regions of DIC, ROBL) were built using PDB-7Z8F^27^ as the starting model. The dynactin filament was built based on PDB-7Z8F and PDB-6ZNL^28^.

The Spindly motif and its flanking residues interacting with the dynactin pointed end complex were built manually guided by the density and the AlphaFold2 prediction of the Spindly motif bound to p25 and p27 dynactin subunits. The longer coiled-coil at the pointed end was assigned to JIP3 LZII as we observed fragmented density connecting the Spindly motif to this coiled-coil. Additionally, LZII is the only sequence in JIP3 that can form a coiled-coil of sufficient length to match our density. The lower short coiled-coil at the pointed end had a similar length as the JIP3 RH2 domain and was therefore assigned to this sequence. The orientation of the RH2 domain was determined based on LZII. AlphaFold2 prediction of JIP3 1-600 was used to obtain the starting models for the LZII and RH2 domains.

The IC-LC tower bound to DHC helical bundles 8/9 was modelled using a combination of the PDB-7Z8F and the AlphaFold2 prediction of IC-LC tower bound to a single DHC. The AlphaFold2 model predicted the interaction of LC8 with DHC tail-1 as can be seen from the fit in the experimental density (Figure S5B). DHC tail-2 was built taking PDB-7Z8F as the starting point followed by flexible fitting in COOT.

The p150 ICD and CC2 were built using an AlphaFold2 model of these segments. The p150 CC1B-IC-LIS1-N AlphaFold2 prediction was used to model CC1B, IC^1–32^ and LIS1-N. The predicted model was docked in the experimental density based on the IC^1–32^, LIS1-N and CC1B. The orientation of the IC^1–32^ helices and LIS1-N along with the spacing between these two binding sites on CC1B were identical in both the AlphaFold2 model and the cryo-EM density. This also aided in identifying the registry of the CC1B fragment. The built registry of the CC1B coiled-coil was further confirmed based on hydrophobic residues 513-520, which could be distinguish at the resolution of ∼4.2 Å. CC1B bends around the LIS1-N, which was built using flexible fitting in COOT followed by refinement in PHENIX.

Dynein-A motor domain was built using PDB-5NUG^14^. The cryo-EM density of dynein-A was consistent with ADP in AAA1, ATP in AAA2 and ADP in AAA3. Nucleotide density in AAA4 was fragmented but correlated well with an ADP which is in line with other structures of the dynein motor domain in pre-powerstroke state (PDB-4RH7^124^ and PDB- 8FDT^83^). Dynein-B motor domain was built using PDB- 7Z8G^27^ which was solved under similar buffer conditions. Cryo-EM density of dynein-B was consistent with ADP in AAA1, ATP in AAA2 and AMP-PNP in AAA3 and AAA4 in line with the previous structure.

For generating the composite map, the locally refined maps (Figure S2) were placed into the consensus map of dynein-dynactin-JIP3^1–560^–LIS1 filtered to 20 Å using UCSF Chimera^123^. The maps were then aligned with respect to each other using overlapping regions using the ‘fit-in-map’ function. The maps were then merged using the ‘volume max’ function and then filtered to 15 Å. Although the raw data show that there is flexibility in the orientation of the complex relative to the MT (Figure S2), for illustrative purposes the composite map was overlaid onto the 13 protofilament microtubule (from^27^) by matching the configuration of the microtubule binding domains of dynein-B with the tubulin dimers with PDB-6RZB^125^.

### AlphaFold2 prediction

All structure predictions were performed using AlphaFold2 through a local installation of ColabFold^126^ running MMseqs2^127^ for homology searches and AlphaFold2^128^ or AlphaFold2-Multimer^129^ for the predictions of single or multiple chains, respectively. JIP3 (Uniprot Q9UPT6) regions 1-185 and JIP3 1-600 were predicted by running ColabFold 1.2.0 on two copies of these segments. The RH1 domain-DLIC^helix^ interaction was predicted by running ColabFold 1.2.0 on two copies each of JIP3 1-99 and DLIC2 427-439 (Uniprot O43237). IC-LC tower was predicted by running ColabFold 1.3.0 using two copies each of DIC (100- 150) (Uniprot Q13409), Tctex1 (Uniprot P63172) and LC8 (Uniprot P63167). Interaction of IC-LC tower with dynein heavy chain was predicted by running ColabFold 1.3.0 using two copies each of DIC (100-150), TCTEX1, LC8 and a single copy of dynein heavy chain (1110-1350) (Uniprot Q14204). Dynactin p150 (Uniprot Q14203-1) regions were predicted by running ColabFold 1.3.0 using two copies of DCTN1 (1-556) for Cap-Gly domain to the CC1B segment and of DCTN1 (557-987) for the ICD-CC2 segment. LIS1 (Uniprot P43034) was predicted by running ColabFold 1.3.0 using two copies of this protein. Interaction between CC1B-DIC-N-LIS1-N was predicted by running ColabFold 1.3.0 on two copies each of DCTN1 (350-547), DIC (1-34) and LIS1 (1-84). The adaptors JIP3, HOOK3 (Uniprot Q86VS8), BICDR1 (Uniprot Q6ZP65), BICD2 (Uniprot Q8TD16), RILP (Uniprot Q96NA2), RILPL1 (Uniprot Q5EBL4) and RILPL2 (Uniprot Q969X0) were predicted by running ColabFold 1.2.0 on two copies of each protein. Interactions of RH1 domains with C-terminal helical elements in RILP, RILPL1 and RILPL2 were predicted by running ColabFold 1.3.0 using two copies of residues 21-103 and 337-353 for RILP, residues 1-141 and 353-367 for RILPL1 and residues 25-106 and 185-211 for RILPL2. The predicted aligned error (PAE) with respect to residue was mapped onto the predicted structure using PointPAE 1.0 (https://github.com/sami-chaaban/PointPAE) (doi: 10.5281/zenodo.6792801), and ChimeraX^130^ was used for visualization.

### Size-exclusion chromatography-multi angle light scattering (SEC-MALS)

SEC-MALS measurements were performed using a Wyatt Heleos II 18 angle light scattering instrument coupled to a Wyatt Optilab rEX online refractive index detector. 70 μL of each sample at 1 mg/mL was fractionated in GF150 buffer using a Superdex 200 5/150 analytical gel filtration column (GE Healthcare) run at 0.1 mL/min on an Agilent 1200 series LC system collecting UV at 280 nm before then passing through the light scattering and refractive index detectors in a standard SEC MALS format. Protein concentration was determined from the excess differential refractive index based on 0.186 ΔRI for 1 g/mL. The measured protein concentration and scattering intensity were used to calculate molecular mass from the intercept of a Debye plot using Zimm’s model as implemented in the Wyatt ASTRA software. The instrumental setup was verified using a 2 mg/mL BSA standard run of the same volume as experimental runs. The BSA monomer peak was used to check mass determination and to evaluate interdetector delay volumes and band broadening parameters that were then applied to experimental runs during analysis.

### Dynein motor domain pulldown

Equilibration buffer (GF150 buffer supplemented with 0.1 mM Mg-ATP and 10% glycerol) was used to equilibrate 20 μL of SNAP-Capture Magnetic Beads (New England Biolabs) slurry in 1.5 mL Protein Lo Bind Tubes (Eppendorf). SNAP-tagged LIS1 constructs were first covalently coupled to beads by incubating 50 μL of 1 μM of the respective protein at room temperature for 30 min. Beads were washed once with 1 mL of equilibration buffer and once with 1 mL binding buffer (GF150 supplemented with 0.05% IGEPAL (MilliporeSigma), 1 mM ATP, 1 mM Na3VO4 (New England Bioscience) and 10% glycerol). Beads with then incubated with 50 μL solution containing 0.2 μM dynein motor domain gently shaken for 1 hr at room temperature. The supernatant was removed and beads were washed twice with 500 uL binding buffer. 40 μL of SDS loading dye (ThermoFisher Scientific) was then added to the beads followed by incubation at 95°C for 5 min. 20 μL of this sample was analysed via SDS-PAGE and visualized by Coomassie Blue staining. The abundance of dynein motor was determined using densitometry in ImageJ. Three technical replicates were done per condition.

### Cell lines

HeLa GFP-BICD2N(1-400)-MTS and GFP-MTS cell lines were generated using the FLP-FRT recombination system. More specifically, Flp-In T-REX HeLa cells were transfected with 1.8 μg of pOGG4 plasmid and 200 ng of pcDNA5/FRT/TO plasmid encoding either a GFP-BICD2N-MTS or a GFP-MTS cassette using FuGENE HD (Promega). Cells with a stably integrated transgene were selected with 150 μg/mL hygromycin over 4-5 days. Cells were maintained in Dulbecco’s modified Eagle’s medium supplemented with 10% FBS and 1% penicillin/streptomycin. Transgene expression was induced by adding 10 μg/mL tetracycline to the culture medium.

### Mitochondrial relocation assay

HeLa GFP-BICD2M-MTS cells were transfected with 20 nM of either Non-Targeting siRNA #1 D-001810-01-05 (Dharmacon), ON-TARGETplus *PAFAH1B1* siRNA J- 010330-07-0002 (Dharmacon) or SMARTPool: ON-TARGETplus *DYNC1H1* siRNA L-006828-00-0005 (Dharmacon) using Lipofectamine RNAiMax (ThermoFisher). Electroporations were performed 48 hours after siRNA transfection using the Neon Transfection System (ThermoFisher). Cells were resuspended in Resuspension Buffer R (ThermoFisher) at a density of 20×10^6^ cells/mL. 13 μL of cells were mixed with 2.5 μL of purified protein for a final protein concentration of 55 nM. Cells were pulsed twice at 1400 V for 2 ms using a Neon pipette tip and were immediately recovered in pre-warmed DMEM medium containing 10% FBS and 10 μg/mL tetracycline. Electroporated cells were plated on an imaging dish pre-coated with poly-D-lysine and incubated at 37˚C for 8 hours to allow for cell spreading. Staining was performed with 20 μM Hoechst (ThermoFisher), 5 μg/mL CellMask Orange (ThermoFisher) and 50 nM MitoTracker Deep Red (ThermoFisher) for 20 minutes at 37˚C. Samples were imaged using a Nikon Ti2 inverted fluorescence microscope with a 20x/0.75NA Air lens.

### Immunoblotting analysis and antibodies

For Western blot analysis, membranes were blocked with 5% milk in TBST for 1 hour at room temperature. Primary antibody incubation was carried out in 5% milk in TBST either for 2 hours at room temperature or for 14 hours at 4˚C. Secondary antibody incubation was performed in 5% milk in TBST for 1 hour at room temperature. Primary antibodies used were: Mouse monoclonal anti-LIS1 (Cat no. sc-374586, Santa Cruz Biotechnology, 1:1000 dilution), mouse monoclonal anti-Dynein IC1/2 (Cat no. sc-13524, Santa Cruz Biotechnology, 1:1000 dilution), Mouse monoclonal anti-GAPDH (Cat no. ab8245, Abcam, 1:1000 dilution), Rabbit polyclonal anti-SNAP-tag (Cat no. P9310S, New England Biolabs, 1:1000 dilution). Secondary antibodies used were: Goat anti-mouse IgG HRP (Cat no. P0447, Dako, 1:1500 dilution) and Goat anti-rabbit IgG HRP (Cat no. ab6721, Abcam, 1:2000 dilution).

### Quantification of mitochondrial spread

The spread of mitochondria relative to the nucleus was quantified using a custom ImageJ script (available at https://github.com/jboulanger/imagej-macro/tree/main/Cell_Organization) as previously described^131^. The mitochondrial regions, the nucleus and the cell contour were defined by segmentation of MitoTracker, Hoechst and CellMask signals, respectively. The mitochondrial spread was measured as the trace of the second moment matrix of the MitoTracker signal intensity in each cell, akin to a standard deviation in 2D.

### Quantification and statistical analysis

Statistical analyses were performed with GraphPad Prism software or in R^132^. The statistical details and specific statistical test used is mentioned in the respective figure caption. Plots were made with GraphPad Prism or in R

## Acknowledgements

We thank S. Chaaban for discussions, assistance in processing microtubule bound dynein/dynactin complexes and AlphaFold2 modelling. We thank S. Bullock, S. Chaaban and E.A. d’Amico for critical reading of the manuscript. We thank the MRC Laboratory of Molecular Biology Electron Microscopy Facility for access and support of electron microscopy sample preparation and data collection; J. Grimmett, T. Darling and I. Clayson for providing scientific computing resources. We thank Chris Batters for assistance with SEC-MALS measurements. This work was supported by Wellcome [210711/Z/18/Z], the Medical Research Council, as part of United Kingdom Research and Innovation (also known as UK Research and Innovation) [MRC file reference number MC_UP_A025_1011], the Fundação para a Ciência e a Tecnologia (FCT)/Ministério da Ciência, Tecnologia e Ensino Superior [PTDC/BIA-CEL/1321/2021], Boehringer Ingelheim Fonds PhD fellowship to G.M. and the EMBO Postdoctoral Fellowship [ALTF 197-2021] to K.S.

## Author contributions

K.S. and C.L. performed protein purification, EM analysis, structure determination and single molecule assays. G.M. performed all cellular experiments. J.B.G and R.G. performed protein purification and pulldown assays. A.P.C. guided the project. K.S., C.L. and A.P.C. prepared the manuscript with input from all authors.

## Competing Interest

The authors declare no competing interests.

## Data Availability

Atomic coordinates and cryo-EM maps have been deposited in the Protein Data Bank (PDB) or Electron Microscopy Data Bank (EMDB), respectively, under accession codes 8PTK and 17873 (composite dynein-dynactin-JIP3-LIS1), 8PR2 and 17832 (dynein tail – JIP3 HBS1), 8PR3 and 17833 (dynein tail – JIP3 RH1 domain), 8PR4 and 17834 (pointed end–JIP3), 8PQY and 17828 (dynein-A motor domain + LIS1), 8PQW and 17826 (dynein-A motor domain-LIS1- CC1B-DIC-N), 8PQZ and 17829 (dynein-A1/A2 bound to LIS1), 8PQV and 17825 (dynein-B motor domain), 8PR0 and 17830 (dynein-A tail + IC-LC tower + p150), 8PR1 and 17831 (dynein-B tail + IC-LC tower), 17835 (consensus map of dynein-dynactin-JIP3^1–185^–LIS1), 17836 (consensus map of dynein-dynactin-JIP3^1–560^–LIS1) and 8PR5 (autoinhibited model of dynactin p150).

All light microscopy data and SDS-PAGE gels used for quantification are available at 10.5281/zenodo.8124243.

## Supplementary Figures

**Figure S1.**
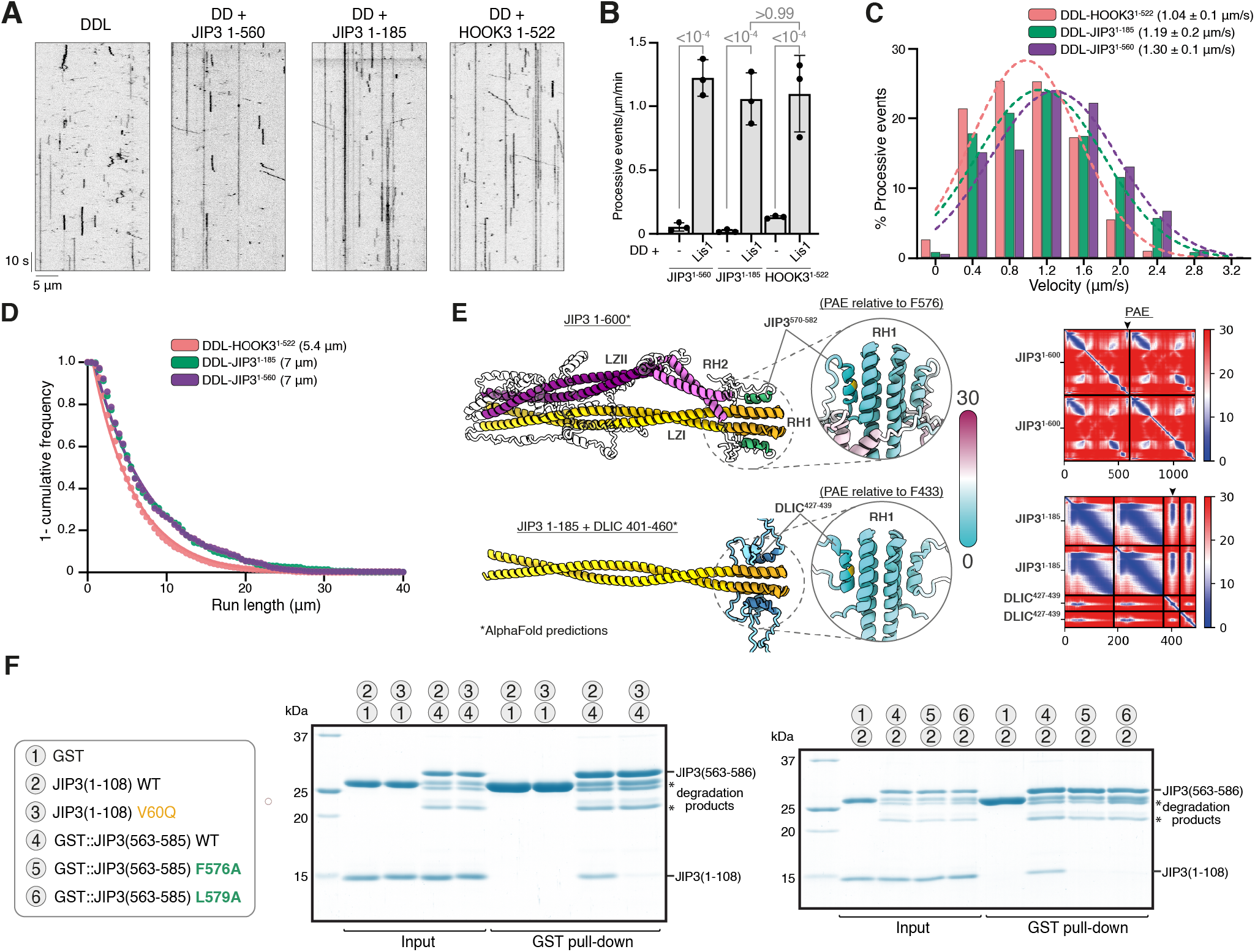
JIP3 is a dynein activating adaptor that undergoes autoinhibition. **(A)** Kymographs of TMR-dynein-dynactin in presence of LIS1, JIP3^1–560^, JIP3^1–185^ and HOOK3^1–522^. **(B)** Quantification of the number of processive events per μm microtubule per minute with the mean ± S.D. plotted. The total number of movements analysed were 26 (DDL), 74 (DD-JIP3^1–560^), 45 (DD-JIP3^1–185^), 224 (DD-HOOK3^1–522^). The data for DD-LIS1-JIP3^1–560^/JIP3^1–185^/HOOK3^1–522^ is the same as displayed in Figure 1C and is shown for comparison purposes. Experiments were performed with three technical replicates and statistical significance was determined using ANOVA with Tukey’s multiple comparison test. **(C)** Distribution of mean velocities of processive events. Gaussian fit of the distribution is shown as dotted lines along with mean velocity ± S.D. (n=3). **(D)** A 1-cumulative frequency distribution plot showing run length for dynein-dynactin-LIS1 with different adaptors fit to a one-phase exponential decay. The decay constant (run length) is shown. **(E)** AlphaFold2 prediction of two copies of JIP3 (1-600) (top) and JIP3 (1-185) along with two copies of DLIC2 (401-460) (bottom). The models are coloured based on PAE values (in Å) relative to that of the highlighted residues (yellow) where lower values represent higher confidence. The full PAE plot is shown on the right. **(F)** Coomassie Blue-stained SDS-PAGE gel of purified recombinant protein mixtures prior to the addition of glutathione agarose resin and of proteins eluted from glutathione agarose resin after GST pull-down. The gel on the left was displayed in Figure 1D after cropping out the lanes for GST controls.

**Figure S2.**
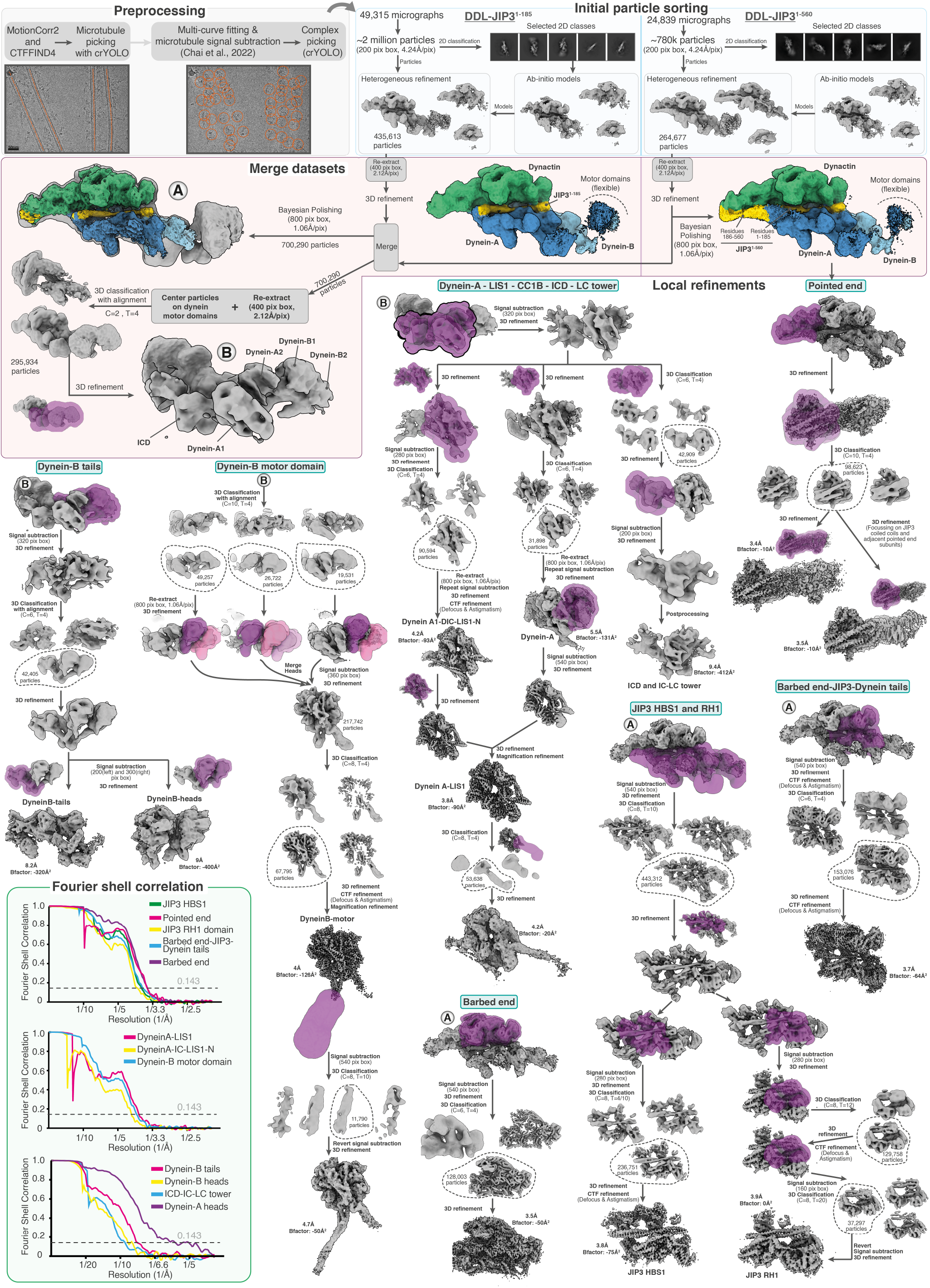
Cryo-EM image processing pipeline for dynein-dynactin-JIP3-LIS1 complexes. (T = Tau fudge, C = number of classes). 3D classifications were performed without alignment unless otherwise specified. All defocus, magnification, and beam-tilt refinements were immediately followed by a 3D refinement (not shown). Plots show the gold standard Fourier shell correlation. The dotted horizontal line shows the 0.143 cut-off.

**Figure S3.**
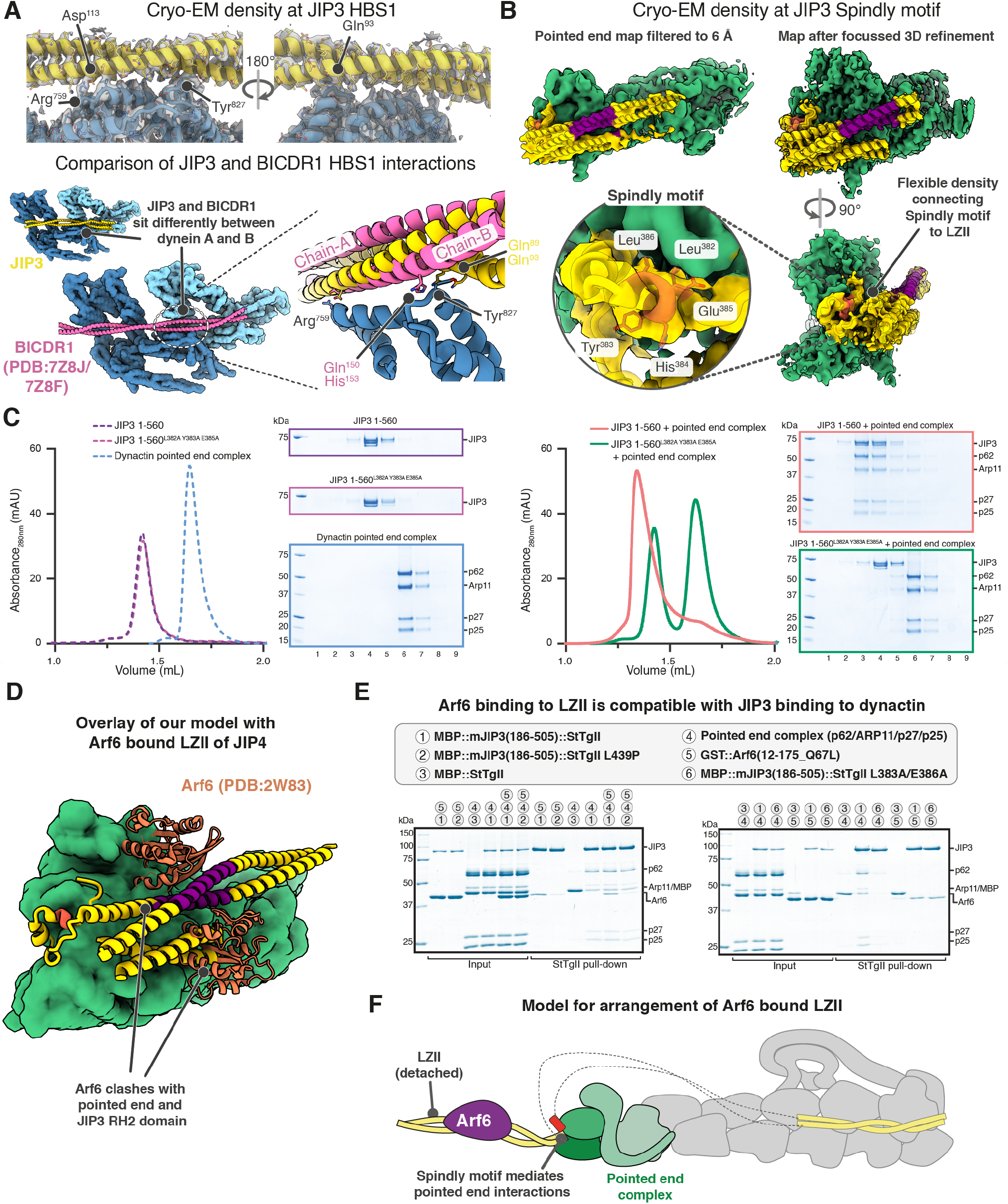
Interactions of JIP3 with dynein-dynactin. **(A)** Cryo-EM density at the JIP3-HBS1 is shown (top). The trajectory of the JIP3 LZI and its interaction with dynein heavy chain are compared to those of BICDR1^27^. **(B)** Cryo-EM density at the dynactin pointed end is shown. **(C)** Size-exclusion chromatography elution profiles from a Superose 6 3.2/300 increase column (left) and Coomassie Blue-stained SDS-PAGE gel (right) to compare complex formation between JIP3^1– 560^ or JIP3^1–560^(L382A, Y383A, E385A) and pointed end complex. **(D)** Overlay of PDB-2W83 (Arf6 bound to JIP4-LZII)^73^ with our model of JIP3 bound at the pointed end. **(E)** Coomassie Blue-stained SDS-PAGE gel of purified recombinant protein mixtures prior to the addition of Strep-Tactin Sepharose resin and of proteins eluted from the resin after strep-tag (StTgII) pull-down. **(F)** Schematic model for LZII orientation with respect to the pointed end when bound to Arf6.

**Figure S4.**
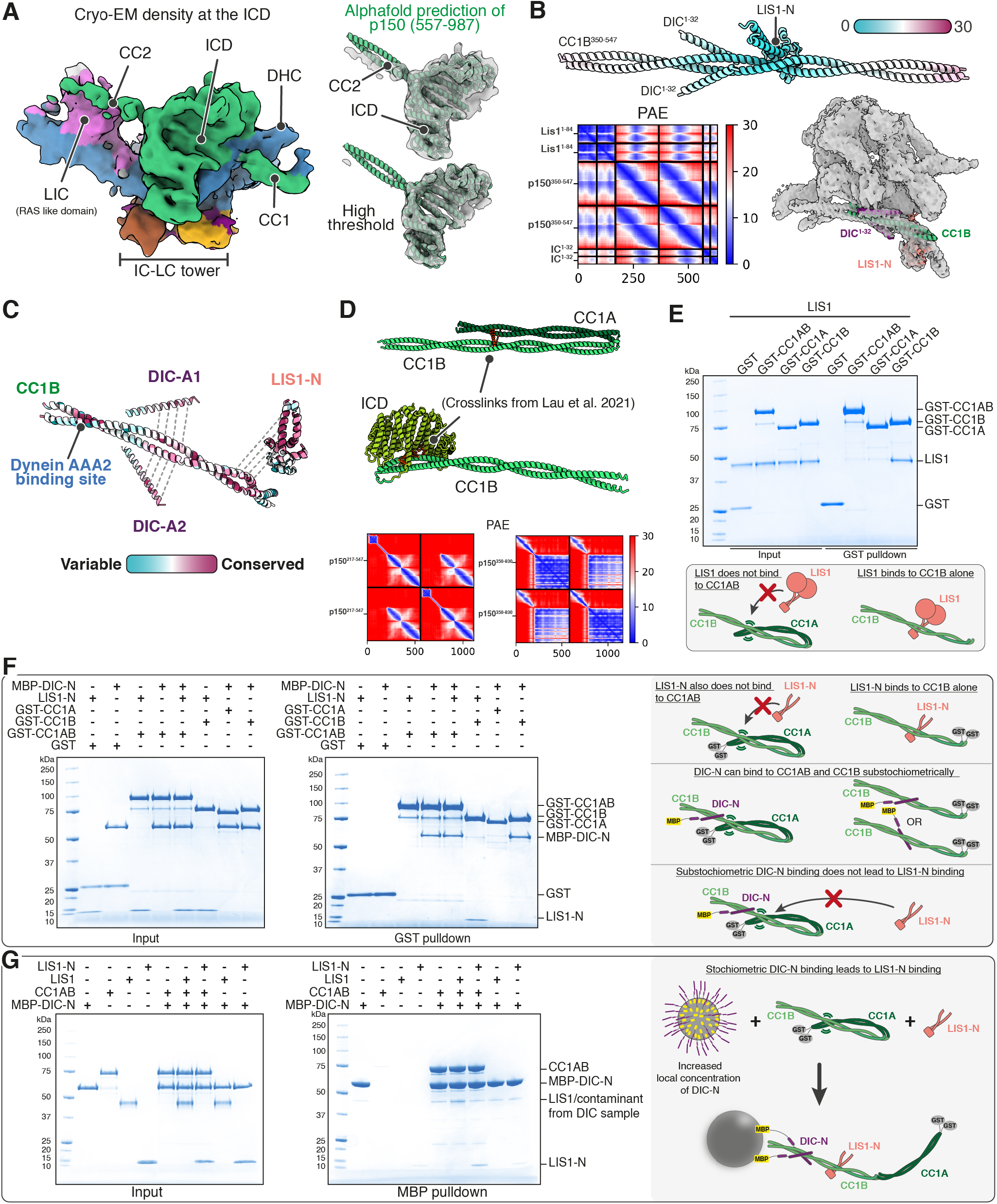
DIC-N helps overcome p150 autoinhibition. **(A)** Cryo-EM map of ICD domain and IC-LC tower bound to dynein-A tail. Fit of AlphaFold2 model of ICD and initial part of CC2 in the cryo-EM map (right) at low threshold (top) and high threshold (bottom). **(B)** AlphaFold2 prediction of DIC^1–32^ and LIS1-N binding to CC1B coloured based on PAE values (in Å) relative to LIS1-K64 (top). The full PAE plot and the fit of this prediction in the experimental density are shown (bottom) **(C)** Cartoon representation of CC1B, DIC^1–32^ and LIS1-N coloured based on sequence conservation determined using the ConSurf web-server^114^ is depicted. **(D)** Arrangement of CC1A/B (top) and CC1B-ICD (middle) predicted by AlphaFold2 is shown. All crosslinks between CC1A-CC1B generated by crosslinking dynactin with BS3^28^ are mapped on the model as red lines. PAE plots corresponding to the predictions are shown (bottom). **(E), (F), (G)** Coomassie Blue-stained SDS-PAGE gels of purified recombinant protein mixtures prior to the addition of glutathione agarose or amylose resin and of proteins eluted from glutathione agarose or amylose resin after pull-down. Schematics summarising the results are depicted next to each experiment.

**Figure S5.**
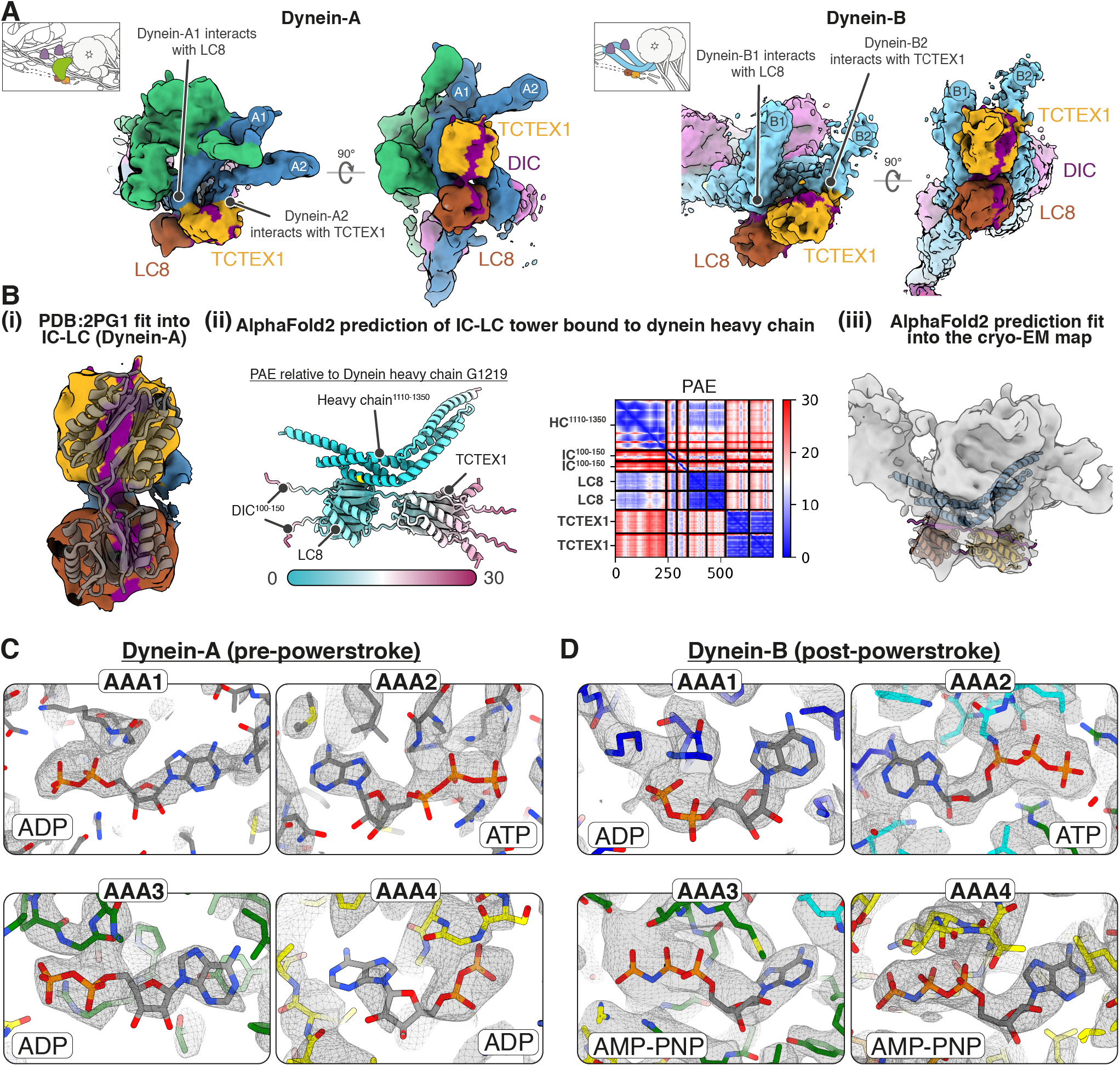
The IC-LC tower bridges the heavy chains of a dynein dimer. **(A)** Cryo-EM maps of IC-LC tower bound to dynein-A (left) and dynein-B (right) heavy chains. **(B)(i)** Crystal structure of IC-LC tower (PDB-2PG1)^81^ fit in the cryo-EM density of dynein-A IC-LC tower. **(ii)** AlphaFold2 model of dynein heavy chain with IC-LC tower coloured based on the relative PAE (in Å) to dynein heavy chain residue G1219 and the full PAE plot are shown (top). The fit of this prediction into the cryo-EM density of dynein-A IC-LC tower is shown **(iii)**. **(C)** Model and density in the nucleotide pocket of AAA1-4 of dynein-A. **(D)** Model and density in the nucleotide pocket of AAA1-4 of dynein-B.

**Figure S6.**
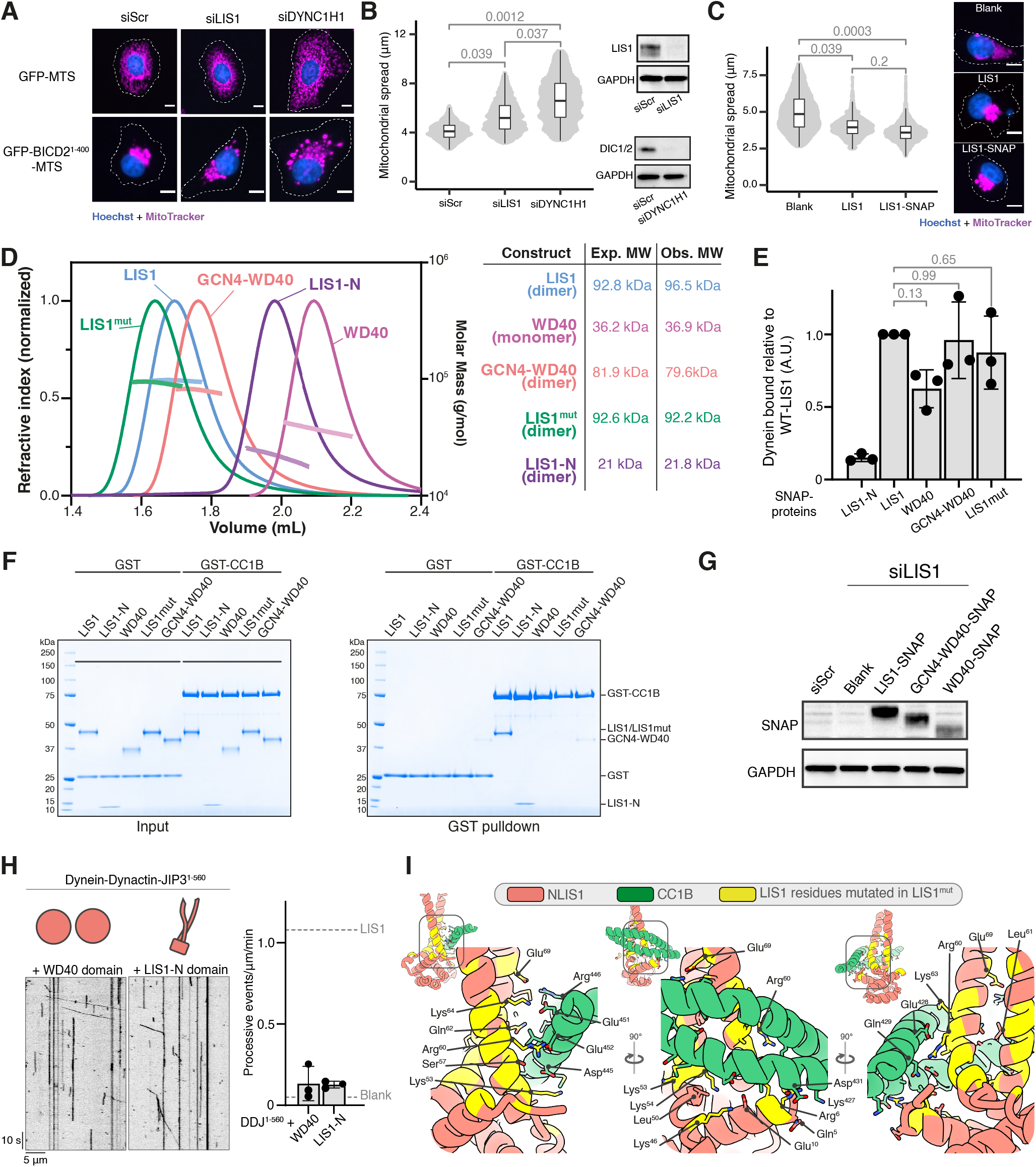
LIS1 requires both LIS1-N and WD40 domains to stimulate complex assembly. **(A)** Representative images and **(B)** quantification showing the distribution of mitochondria (magenta) in HeLa GFP-BICD2N-MTS and GFP-MTS cells transfected with siRNAs against LIS1 and the dynein heavy chain (DYNC1H1). A scrambled siRNA was used as a negative control. Scale bars represent 10 μm. Data are plotted from 3 biological replicates. Knockdown efficiency was determined by western blot. Depletion of the dynein intermediate chain (DIC1/2) was used as a proxy for dynein heavy chain knockdown efficiency. **(C)** Comparison of LIS1 knockdown HeLa GFP-BICD2N-MTS cells rescued with either untagged LIS1 or LIS1- SNAP. Scale bars represent 10 μm. Data are plotted from 3 biological replicates. **(D)** SEC-MALS of purified LIS1 constructs. Mean observed molar mass (Obs.) and expected (Exp.) molar mass is indicated. **(E)** Quantification of pulldown of dynein motor domain by SNAP-tagged LIS1 constructs. The data is plotted from 3 technical replicates where the dynein binding was normalized using the LIS1-SNAP condition in each replicate. **(F)** Coomassie Blue-stained SDS-PAGE gel of purified recombinant protein mixtures prior to the addition of glutathione agarose resin and of proteins eluted from glutathione agarose resin after GST pull-down. **(G)** Transfection efficiency of SNAP-tagged LIS1 constructs used for rescuing LIS1 knockdown in Figure 6 was determined by Western blot. **(H)** Kymographs and quantification of processive events per μm microtubule per minute of TMR-dynein-dynactin-JIP3^1–560^ in the presence of (from left) WD40 domains and LIS1-N. Cartoons depicting the LIS1 construct used are shown above each kymograph. Experiments were performed with three technical replicates**. (I)** Amino acids present in the interface between LIS1-N and CC1B. The amino acids coloured in yellow were mutated to generate the LIS1^mut^ construct used in Figure 6D and E. All statistical significance values were determined using ANOVA with Tukey’s multiple comparison.

**Figure S7.**
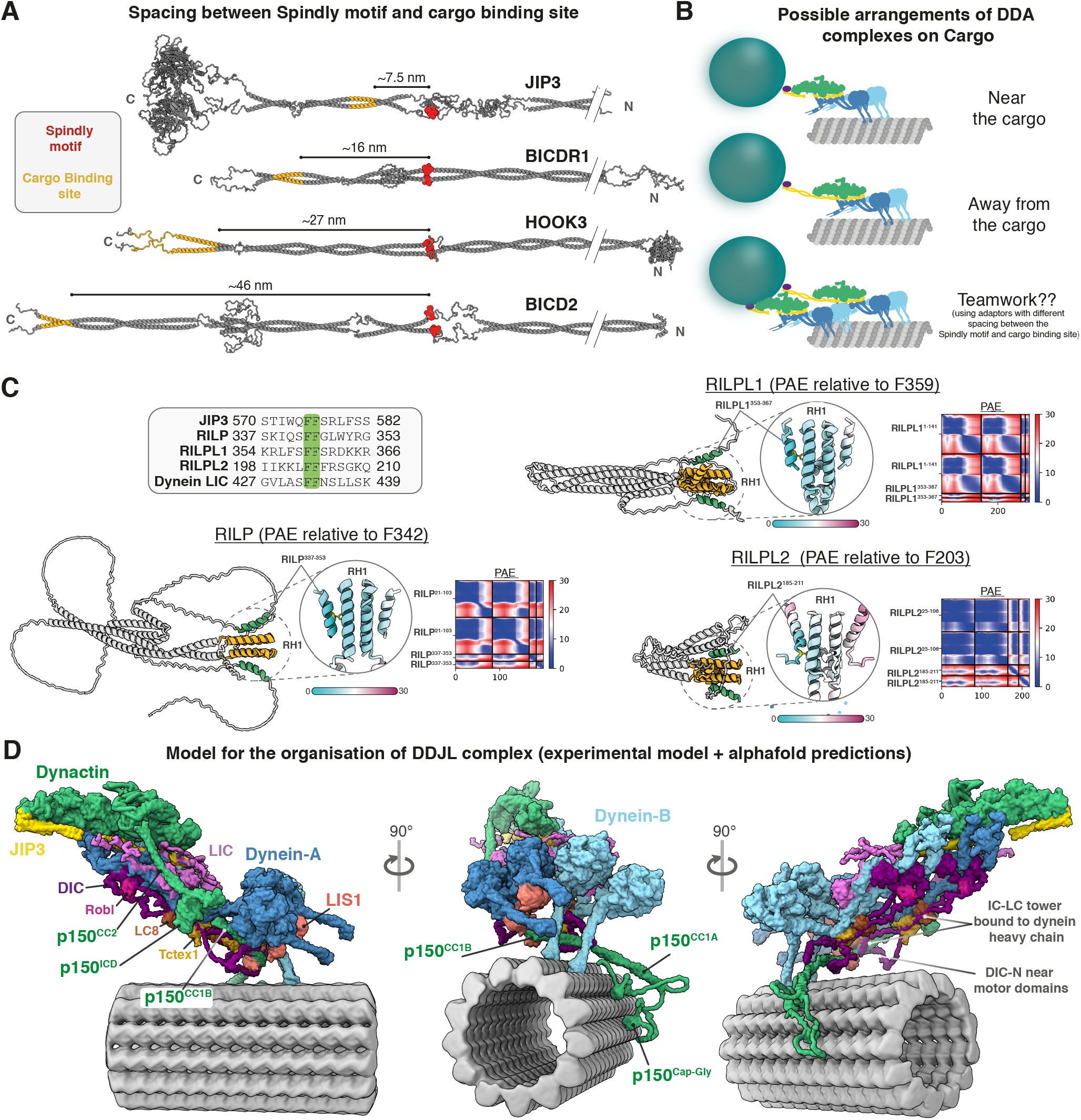
Model for organisation of DDJL complex on microtubules. **(A)** Linearized AlphaFold2 models of the adaptors JIP3, BICDR1, HOOK3 and BICD2. The Spindly motif (red) and cargo binding regions (yellow) are mapped onto the model to show the approximate distance between the two sites. **(B)** Schematic showing possible arrangements of DDA complexes when recruited by different adaptors. **(C)** AlphaFold2 prediction of RH1 domain-containing proteins (RILP, RILPL1 and RILPL2). These proteins contain a C-terminally located helix containing phenylalanine residues (highlighted in green) that are predicted to bind the RH1 domain in a similar manner as JIP3. **(D)** Model for the overall organisation of the DDA-LIS1 complex on microtubules. The N-terminal segments of p150 until CC1A and unstructured regions of DIC and DLIC were too flexible to be visualized in our cryo-EM structure and have been placed manually to illustrate where these segments are likely to be located in the complex.

## Supplementary Tables

**Table S1. Quantification and statistics of *in vitro* and cellular assays.**

**Table S2. Cryo-EM data collection and refinement statistics**

## Notes

### Competing Interest Statement

The authors have declared no competing interest.

### Summary of Updates

This version of the manuscript is a significant update, providing new insights not only into dynein activation by JIP3, but also revealing the intricate interactions between dynein and dynactin during the formation of an active complex and how LIS1 stimulates this process.

